# Distinct cancer-associated fibroblast states drive clinical outcomes in high-grade serous ovarian cancer and are regulated by TCF21

**DOI:** 10.1101/519728

**Authors:** Ali Hussain, Veronique Voisin, Stephanie Poon, Jalna Meens, Julia Dmytryshyn, Victor W Ho, Kwan Ho Tang, Joshua Paterson, Blaise Clarke, Marcus Q Bernardini, Gary D Bader, Benjamin G Neel, Laurie E Ailles

**Affiliations:** Department of Medical Biophysics, University of Toronto, Toronto, ON, Canada; The Donnelly Centre, University of Toronto, Toronto, ON, Canada; Princess Margaret Cancer Centre, University Health Network, Toronto, ON, Canada; Laura and Isaac Perlmutter Cancer Center, New York Langone Health, New York, NY; Department of Laboratory Medicine and Pathobiology, University of Toronto, Toronto, ON, Canada; Department of Pathology, University Health Network, Toronto, ON, Canada; Division of Gynaecologic Oncology, University Health Network, Toronto, Ontario, Canada; Department of Molecular Genetics, University of Toronto, Toronto, ON, Canada

**Author notes:** To whom correspondence should be addressed: Laurie Ailles, Princess Margaret Cancer Centre, 101 College St Rm 12-305, Toronto, ON, CANADA, M5G 1L7.

## Abstract

Recent studies indicate that cancer-associated fibroblasts (CAFs) are phenotypically and functionally heterogeneous. However, little is known about CAF subtypes, the roles they play in cancer progression, and molecular mediators of the CAF “state”. Here we identify a novel cell surface pan-CAF marker, CD49e, and demonstrate that two distinct CAF states, distinguished by expression of fibroblast activation protein (FAP), co-exist within the CD49e+ CAF compartment in high grade serous ovarian cancers. We show for the first time that CAF state influences patient outcomes, and that this is mediated by the ability FAP-high (FH) but not FAP-low (FL) CAFs to aggressively promote proliferation, invasion and therapy resistance of cancer cells. Overexpression of the FL-specific transcription factor TCF21 in FH CAFs decreases their ability to promote invasion, chemoresistance and *in vivo* tumor growth, indicating that it acts as a master regulator of the CAF state. Understanding CAF states in more detail could lead to better patient stratification and novel therapeutic strategies.

**One sentence summary:** In this study we demonstrate that cancer-associated fibroblasts (CAFs) in high-grade serous ovarian cancer are heterogeneous, that CAF state drives cancer aggressiveness and patient outcomes, and that TCF21 is a master regulator of CAF state.

## Introduction

High grade serous ovarian cancer (HGSOC) is the most common histological subtype of ovarian cancer and is typically diagnosed at an advanced stage (*1*). Optimal surgical debulking and platinum/taxane-based chemotherapy significantly increase the survival of HGSOC patients, but the vast majority relapse within 5 years of diagnosis and die of their disease (*1*). Due to early dissemination and implantation of cancer cells within the peritoneal cavity, HGSOC patients typically present at late stage with widespread abdominal disease, and nearly invariably develop chemotherapy resistance. In spite of recent advances with targeted therapies such as PARP inhibitors (*2*), bevacizumab (*3*), and immune checkpoint blockade (*4*), these approaches do not currently benefit all patients and mortality rates remain high. The development of more effective treatments for HGSOC patients thus remains a necessary and important goal.

Cancer-associated fibroblasts (CAFs) are a key component of the tumor microenvironment and have several differences relative to their normal counterparts, including increased proliferation, extracellular matrix production, and expression of cytokines and growth factors (*5*). In many cancers, including HGSOC, CAFs have important effects on tumor behavior, including defining the rate and extent of cancer progression through inhibition of cancer cell apoptosis, induction of cancer cell proliferation, promotion of cancer cell migration and invasion and mediation of chemotherapy resistance (*6-10*). More recently CAFs have also been shown to mediate immune-suppression (*11-13*), adding another layer of complexity to their pro-tumorigenic role. A variety of markers have been used to identify CAFs, including α-smooth muscle actin (SMA), platelet-derived growth factor receptors (PDGFRs) and fibroblast activation protein (FAP), and most studies have focused on CAFs that express these markers. More recent studies have shown that CAFs are heterogeneous, and CAF subtypes with distinct phenotypes have begun to be identified in various malignancies (*14-18*). However, while CAFs with distinct phenotypes have been identified in several cancers, the functional characterization of these cells and their effects on tumor progression and patient outcomes have not yet been revealed. Furthermore, molecular mechanisms driving epigenetic differences between CAF subtypes remain uncharacterized.

Here we describe the identification of CD49e as a novel cell surface marker for fibroblasts within HGSOC primary tumor tissues, and discover two distinct CAF states that exist within the CD49e+ fibroblast compartment and can be distinguished based on fibroblast-activation protein (FAP) expression. We demonstrate that FAP-high (FH) and FAP-low (FL) CAFs co-exist at varying ratios in individual tumors and, importantly, CAF status drives patient outcomes. Purified FH and FL CAFs have distinct transcriptional signatures that are prognostic in the TCGA cohort, and i*n vitro* and *in vivo* functional assays reveal differences in their ability to promote cancer cell proliferation, invasion and chemo-resistance. Finally, we show that transcription factor TCF21 is a master regulator of the CAF state. Our extensive molecular and functional characterization of CAFs and analysis of CAF-derived gene signatures in relation to patient outcomes provides novel insights into the significant role of this cell population in HGSOC disease progression, and the potential of manipulating the CAF state as a therapeutic strategy.

## Results

### Isolation and transcriptional profiling of CAFs from primary HGSOC tumor samples

All tumor samples used in this study are listed in Table S1a. Purification of viable CAFs directly from primary tumors using fluorescence activated cell sorting (FACS) requires a robust cell surface marker. PDGFR-β and FAP are commonly used CAF markers but we found these to be either dimly or inconsistently expressed in single cell suspensions from primary HGSOC samples, in line with other studies showing that expression of established CAF markers is heterogeneous and non-overlapping (*18, 19*). We therefore used high-throughput flow cytometry with a panel of 363 antibodies targeting cell surface proteins (*20*) to analyze cultured CAF lines derived from four HGSOC patients and single cell suspensions from five primary HGSOC samples. The latter were co-stained for CD45 and CD31 to allow exclusion of contaminating immune and endothelial cells, respectively. From this screen we identified several proteins that were uniformly highly expressed on the cultured CAFs, but only stained a minority of the CD45-/CD31-cells from primary HGSOC samples, which would be expected to contain a mixture of cancer cells and CAFs (**Figure S1**). The greatest difference was seen for CD49e (ITGA5), which was selected for follow-up. Immunofluorescence studies of HGSOC sections confirmed that CD49e antibody selectively stained the tumor stromal compartment, while pan-CK antibody, as expected, stained tumor cells (**Figure 1A**). Co-staining with an anti-EpCAM antibody (an epithelial cell marker) enabled clear distinction and isolation of CAFs from HGSOC samples as the CD45-CD31-EpCAM-CD49e+ fraction, which varied in frequency between patients (**Figure 1B**). The identity of isolated CD45-CD31-EpCAM+CD49e-(referred to as EpCAM+) and CD45-CD31-EpCAM-CD49e+ (referred to as CD49e+) fractions as cancer cells and CAFs, respectively, was further validated by generating cytospins of the purified populations and staining them for pan-CK, Vimentin and p53. The CD49e+ fraction was positive for Vimentin and negative for pan-CK, as would be expected for a fibroblast population (**Figure 1C**). In addition, in patients for whom the EpCAM+ fraction showed strong nuclear P53 staining, indicative of mutant P53 (*21*), the CD49e+ fraction was negative for P53 staining, demonstrating that CD49e+ cells do not bear the cancer-associated mutation and ruling out the possibility that they are cancer cells that underwent an epithelial-to-mesenchymal transition (EMT; **Figure 1D**).

**Figure 1.**
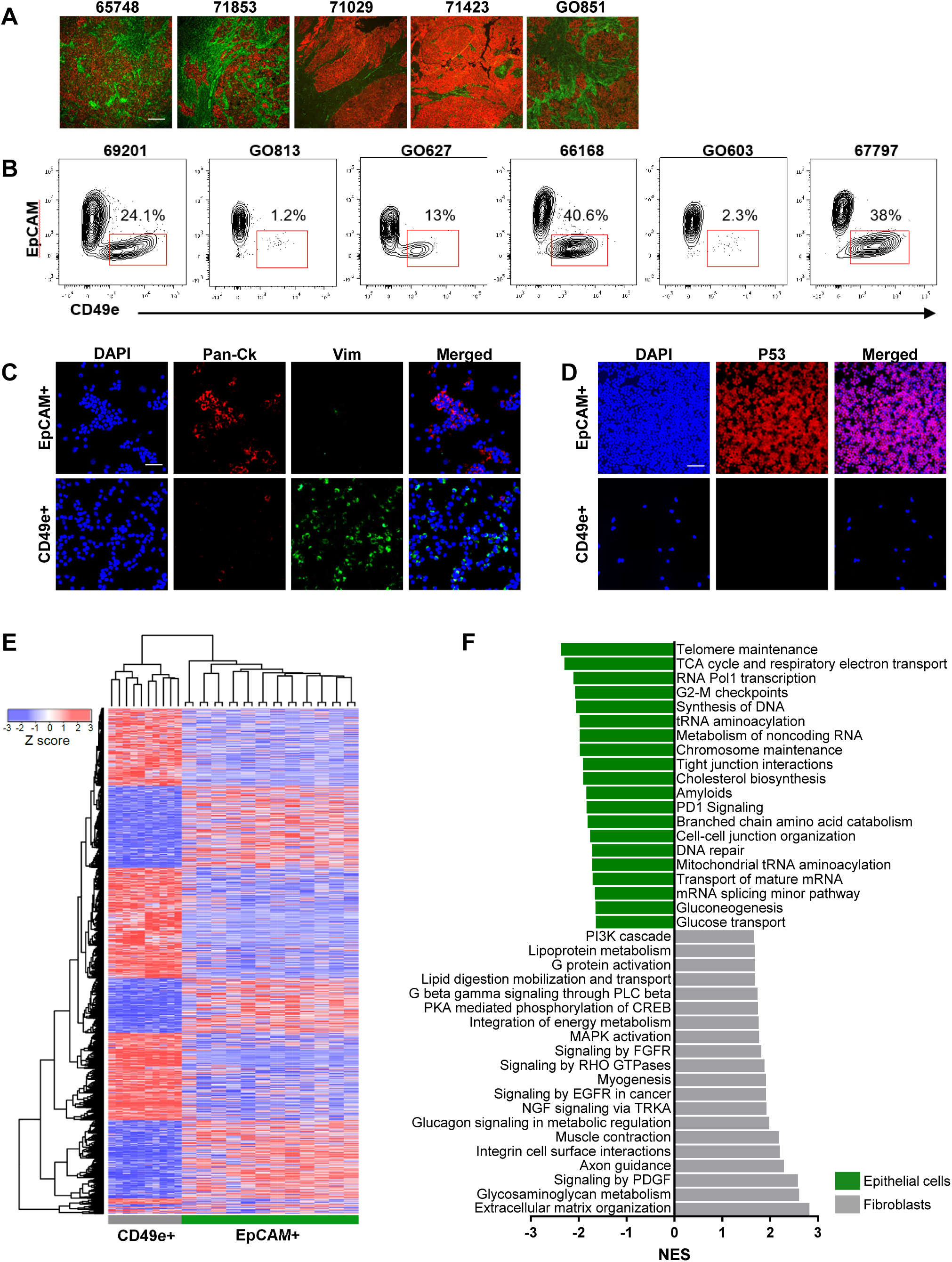
Prospective isolation and transcriptional profiling of CAFs from primary HGSOC samples. **(A)** Representative images of immunofluorescence on 5 different HGSOC samples using pan-cytokeratin (CK; red) and CD49e (green) antibodies. Scale bar = 100 µm. **(B)** Representative FACS plots from 6 different patients showing the viable, CD45-CD31-population stained for EpCAM and CD49e. **(C)** Representative images of cytospins made from isolated EpCAM+ or CD49e+ cells stained for pan-CK (red), Vimentin (green), and Hoechst (blue). Scale bar = 50 µm. **(D)** Representative images of cytospins made from isolated EpCAM+ or CD49e+ cells stained for p53 (red) and Hoechst (blue). Scale bar = 50 µm. **(E)** Heatmap showing unsupervised hierarchical clustering of CD49e+ and EpCAM+ cells isolated directly from HGSOC tumor specimens and profiled by Illumina HT-12v4 microarrays. The EpCAM+ cells were separated into CD133+ and CD133-fractions. The heatmap shows EpCAM+ populations from 12 patients and CD49e+ samples from 10 patients due to data being of poor quality from two of the CD49e+ samples. **(F)** Gene set enrichment analysis of the genes differentially expressed in the CD49e+ *vs* EpCAM+ populations. The top 20 non-redundant gene sets for each population are shown..

To interrogate the transcriptional profiles of cells isolated directly from primary tumor specimens, FACS was used to isolate the CD49e+ fraction and the EpCAM+ fraction from 12 primary HGSOC samples (**Figure S2A**). The latter was further fractionated into EpCAM+CD133+ and EpCAM+CD133-fractions, as we had previously shown that CD133 was a marker of tumor-initiating cells in the majority of primary HGSOC solid tumors (*22*). RNA was extracted and analyzed using Illumina HT-12v4 microarrays. Upon unsupervised analysis, CD133+ and CD133-subsets had very similar transcriptional profiles and clustered together, whereas the CD49e+ population formed a distinct cluster (**Figure 1E**). The CD49e+ fraction expressed known CAF-associated genes, including *ACTA2, FAP, VIM, POSTN, SPARC*, and multiple collagen genes, among others (**Table S2A**). Gene set enrichment analysis (GSEA) was carried out on differentially expressed genes between CD49e+ cells and EpCAM+ cells using the MsigDB Reactome database. EpCAM+ cells were enriched for genes related to cell proliferation, metabolism and epithelial identity (e.g., tight junction interactions, cell-cell junction organization), whereas the CD49e+ subset was enriched for gene sets involved in extracellular matrix organization, myogenesis and known mesenchymal signaling pathways, such as PDGF, FGF and RHO GTPase signaling (*23, 24*). These results further confirm the identity of our purified CD49e+ population as fibroblasts (**Figure 1F, Table S2B)**.

### The CD49e+ fraction separates into 2 clusters

Unsupervised hierarchical clustering of the gene expression data indicated that the CD49e+ CAF population segregated into two sub-clusters (**Figure 1E, Figure S2B**), suggesting that CAFs derived from HGSOC patients are heterogeneous. One cluster expressed classical CAF-related genes, such as *FAP, TGFβ, COL11A1, SULF1* and inflammatory cytokines (e.g., *IL-6, CXCL12*), among others, whereas the other exhibited a distinct gene expression profile that included low expression of the classical CAF marker *FAP* (**Figure 2A, Table S3A**). We therefore refer to these two groups of patients as “FAP-high” (FH) and “FAP-low” (FL). Several of the FH (*FAP, COL11A1, SULF1*)-and FL (*DLK1, TCF21, COLEC11)-*specific genes were validated by reverse-transcription and quantitative polymerase chain reaction (qRT-PCR) on RNA isolated from the CD49e+ fraction of six patients included in the microarray analysis, three from the FH group and three from the FL group (**Figure 2B**). To test whether the two CAF subtypes were a general feature of ovarian cancer stroma, we analyzed the gene expression data set generated by Leung and colleagues (*25*) who used laser capture microdissection to isolate stromal and epithelial components from 31 HGSOC specimens and 8 normal ovary specimens. We interrogated the expression of the top 500 differentially expressed genes between our FH and FL patients in the stromal samples from this cohort and again found two major clusters (**Figure 2A**). This result verified the ability of our FH *vs* FL gene list to segregate HGSOC patients into two subtypes based on expression of these genes in their stroma. Importantly, the transcriptional profile of normal ovarian stroma was distinct from the FL cancer stroma, indicating that FL CAFs represent a distinct phenotype of stromal cells within HGSOC, and not normal fibroblasts (**Figure S2C-D and Table S3B**).

**Figure 2.**
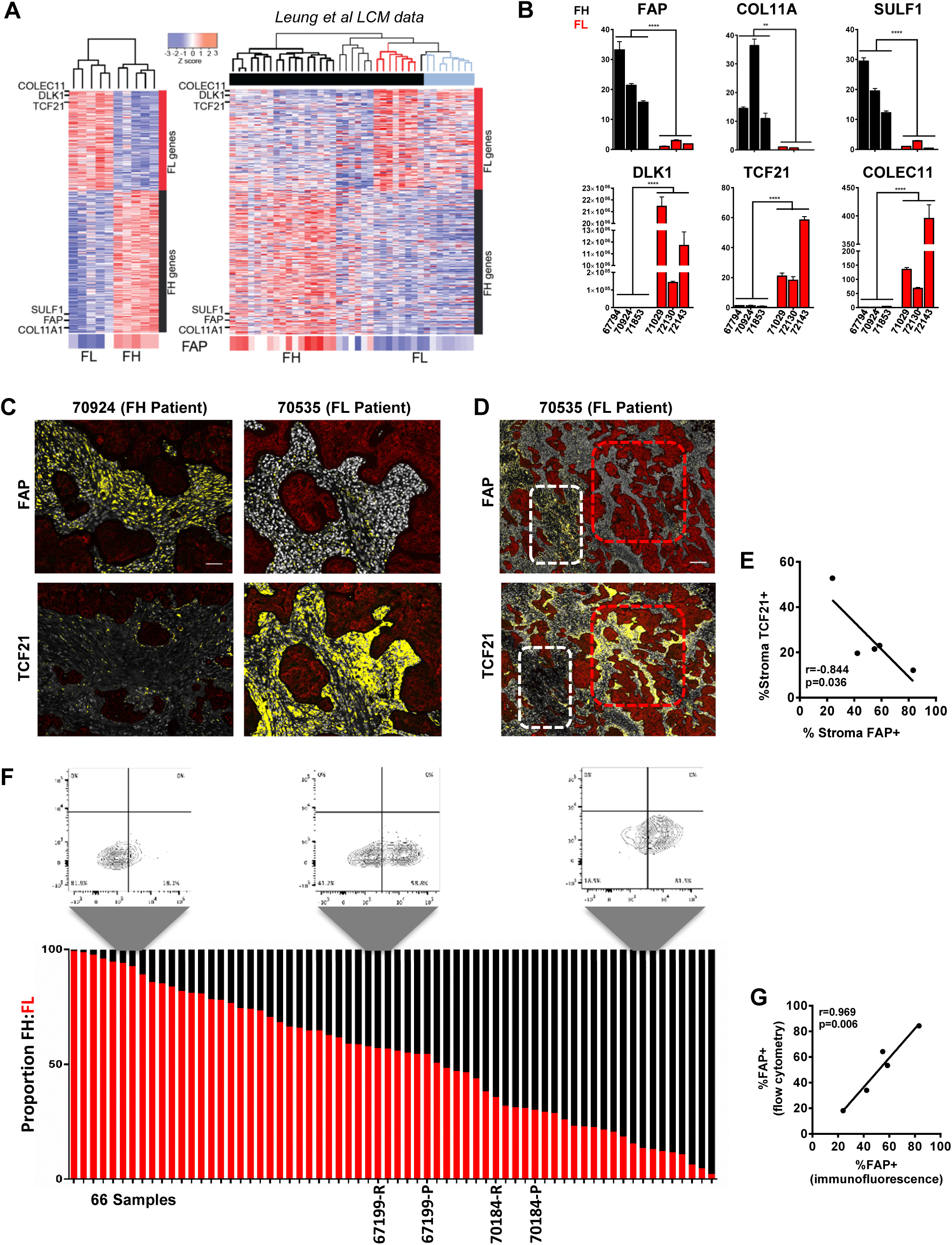
Identification of FH and FL CAFs in HGSOC. **(A)** Heatmap of the top 500 FH vs FL differentially expressed genes (left), and analysis of the same gene list in the Leung *et al* HGSOC and normal ovary stromal samples (right). Black bar: tumor stroma; Blue bar: normal stroma. Black dendrogram: FH patients; Red dendrogram: FL patients. *FAP* gene expression is enlarged at the bottom of the heatmap. **(B)** RNA from the CD49e+ fraction of 3 FH and 3 FL tumors (as classified by microarray) was analyzed by qRT-PCR for the expression of 3 FH-specific and 3 FL-specific genes. Mean±SEM, n=3. *p<0.05; **p<0.01; ****p<0.0001, Student’s t-test. **(C)** Representative images of HGSOC serial sections stained for FAP or TCF21. Sections were co-stained for pan-CK and Hoechst. HALO image analysis software was used to quantify antibody staining within the pan-CK negative stromal regions. Images show stained sections with HALO analysis mask. Red: epithelial cells; White: nuclei within the stromal region; Yellow: FAP or TCF21 positive cells. Scale bar = 50 µm. See Supplemental Figure S3 for additional images. **(D)** Serial sections from patient 70535 stained for FAP or TCF21. White: FAP-high/TCF21-low region; Red: FAP-low/TCF21-high region. Scale bar = 200 µm. **(E)** Graph comparing the % stromal area positive for FAP and TCF21 within the entire tissue section, for 5 tumors analyzed. **(F)** Quantification of the proportion of FH cells within the viable CD45-CD31-EpCAM-CD49e+ fraction by flow cytometry in 64 primary samples and 2 matched recurrences. 3 representative FACS plots, gated on the CD45-CD31-EpCAM-CD49e+ cells and showing FAP staining, are shown above the graph. The two matched primary (P) and recurrent (R) samples are indicated below the graph. **(G)** Graph comparing FAP quantification data obtained using flow cytometry and IF for samples that were analyzed using both methods.

### FH and FL fibroblasts co-exist in the majority of patients

To further interrogate the FH and FL status of patients classified into these groups by gene expression analysis, we performed immunofluorescence (IF) staining for FAP, TCF21 (a highly-expressed FL gene) and pan-CK on serial sections from five tumors categorized as FH (n=3) or FL (n=2) by microarray. The slides were scanned to generate high resolution digital images of the entire section, and the FAP-positive or TCF21-positive stained areas within the Pan-CK negative, stromal regions were quantified using HALO image analysis software (**Figure 2C and Figure S3**). This analysis showed that FH patients indeed had more FAP-expressing cells in their tumor stroma than FL patients, whereas TCF21-expressing cells were more abundant in FL patients. However, we also noticed that within tumor samples, FAP-positive and TCF21-positive cells could both be observed, and regions with predominantly one type of CAF or the other could be identified (**Figure 2D**). Nevertheless, quantification by HALO indicated an anti-correlation between FAP and TCF21 expression (**Figure 2E**). Thus FH and FL CAFs are two distinct subtypes that co-exist within HGSOC tumors at varying ratios.

To more accurately quantify FH and FL CAFs within patient specimens, we performed flow cytometry on 66 HGSOC samples, with the addition of a FAP antibody to the previous combination of CD45, CD31, EpCAM and CD49e, allowing us to quantify FAP expression within the CD49e+ fraction. The FH subset ranged from 0.6% to 98% within the CD49e+ population (**Figure 2F**). Of the 5 patient samples that were also stained by IF, the flow cytometry and immunofluorescence assays showed a strong positive correlation in the proportion of cells expressing FAP (r=0.969).

### Transcriptional profiling of purified FH and FL cells

Our initial gene expression analysis was performed on the bulk CD49e+ fraction, which contained mixtures of FH and FL CAFs at varying ratios, suggesting that the observed clustering of patients into FH and FL groups reflected the predominant population, but these populations were not pure. We therefore carried out RNA-Seq analysis on FH and FL CAFs purified by FACS. For some of these samples, the CD49e+ fraction was predominantly FH or FL, making isolation of both fractions in sufficient numbers difficult; however, for three patients (72143, 70535 and 71423), we successfully generated high-quality RNA-Seq data on both fractions. Principal component analysis of the five FH populations and four FL populations showed that regardless of whether the samples were derived from a FH patient or a FL patient, the samples clustered based on FAP status (**Figure 3A**). Gene set enrichment analysis (GSEA) comparing the genes differentially expressed in isolated FH and FL cells showed a very high concordance with the previously obtained microarray-derived gene lists (**Figure 3B**). In addition, *FAP, COL11A1* and *SULF1* were highly expressed in the FH subset, and *TCF21, COLEC11* and *DLK1* were highly expressed in the FL subset (**Figure 3C**). These results confirm that HGSOC samples contain mixtures of FH and FL CAFs at varying ratios, and that the clustering of patients based on bulk fibroblast profiling was driven by whichever population was dominant in those tumors.

**Figure 3.**
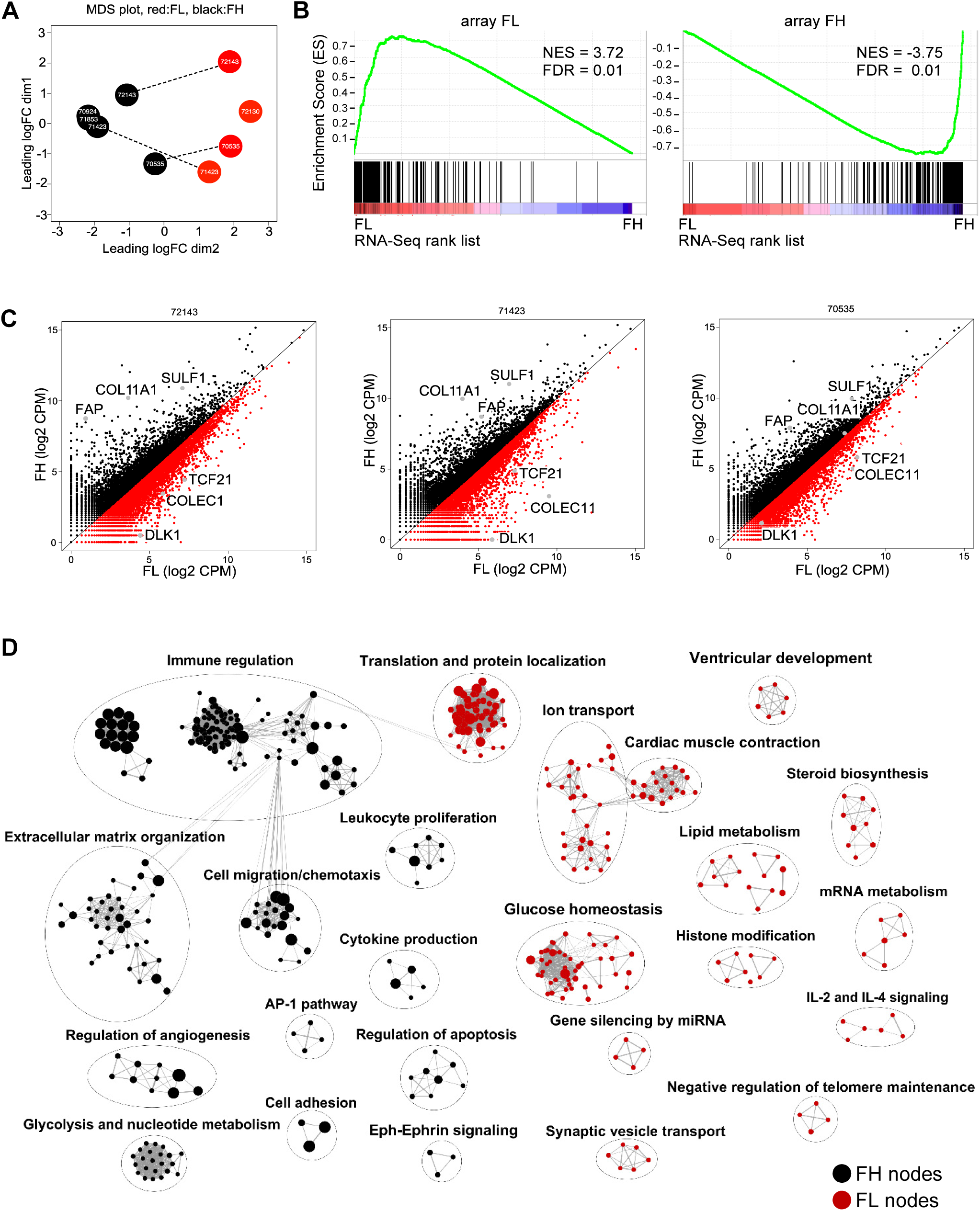
Transcriptional profiling of purified FH and FL CAFs. **(A)** Principal components analysis of RNA-Seq data from FACS-purified FH and FL cells. Dotted lines indicate cases for which FH and FL cells were purified from the same tumor specimen. **(B)** Gene set enrichment analysis of the FH (left) and FL (right) gene lists generated from RNA-Seq data in comparison to the gene lists previously generated by microarrays. **(C)** Scatter plots of FH *vs* FL gene expression in the 3 patients where both populations were isolated from the same patient. The three FH and three FL genes previously used for validation are indicated as gray dots. **(D)** Cytoscape map of the most significantly differentially expressed gene ontology (GO) terms and pathways between FH and FL CAFs. GSEA was performed and the top 300 most differentially expressed GO terms and Reactome pathways are shown. The size of individual nodes correlates with the number of genes in each.

There were 800 differentially expressed genes between FH and FL CAFs at a false discovery rate (FDR) value of ≤0.05 **(Table S4A).** GSEA of the differentially expressed genes between FH and FL cells (**Figure 3D**) showed that FH cells express genes involved in ECM reorganization, cell migration and chemotaxis, immune regulation (including neutrophil activation, regulation of defense response and antigen processing), and regulation of angiogenesis, and thus resemble the classical phenotype that is commonly associated with CAFs in the literature (*11, 23, 26*). By contrast, the most dominant gene sets expressed in FL CAFs include glucose/insulin homeostasis, cardiac muscle contraction and ion transport, translation and protein localization, and lipid metabolism. FL CAFs thus represent a previously unrecognized CAF subtype with a distinct gene expression profile from FH CAFs.

### FH and FL patients can be identified in The Cancer Genome Atlas (TCGA) HGSOC dataset and have distinct clinical outcomes

Gene expression profiling of 489 HGSOC samples by TCGA previously led to identification of four molecular subtypes: mesenchymal, proliferative, differentiated, immunoreactive (*27*). Refined signatures for these subtypes were subsequently generated, and the mesenchymal subtype was found to be associated with worse outcome (*28*). We generated a gene signature based on FH and FL CAF genes by filtering for genes that were both differentially expressed in purified FH *vs* FL cells, and differentially expressed in CAFs compared to EpCAM+ cells in our original microarray data set. This analysis resulted in a list of 165 FH-specific genes and 78 FL-specific genes (Figure S4A, Table S5). We then interrogated this signature against TCGA RNA-Seq data (n = 374 patients) and found that FH and FL transcripts were detectable in a large number of patients, with distinct FH and FL clusters present (Figure 4A). A scoring system was established to classify patients (Figure S4B). When a patient expressed 75% of the FH genes at a level higher than the population mean, the patient was classified FH (shown in black in Figure 4A). A second, less stringent threshold classified patients as FH if they expressed 50% of the FH genes above the mean (shown in grey in Figure 4A). The same rule was applied to classify the FL patients (shown in red and pink, respectively in Figure 4A). The majority of the FH patients overlapped with the TCGA mesenchymal subtype, whereas the FL patients were distributed amongst the mesenchymal, proliferative and differentiated subtypes (Figure 4A, Figure S4C). GSEA shows that the FH genes were highly enriched in the mesenchymal signature (*28*) whereas the FL genes were not (Figure 4B), suggesting that that the mesenchymal subtype is largely driven by the presence of FH fibroblasts.

**Figure 4.**
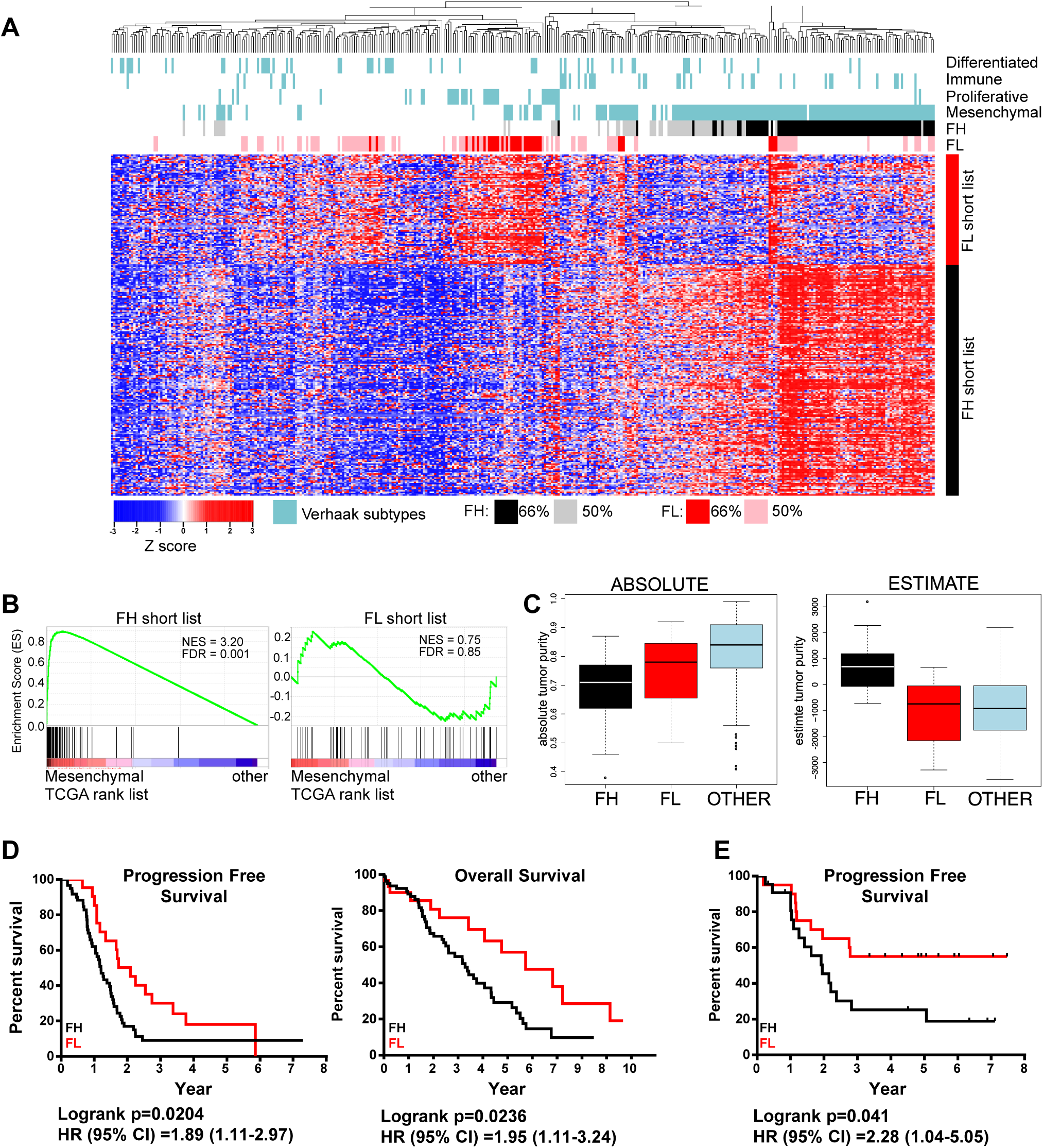
FH patients correspond to the mesenchymal molecular subtype and have worse clinical outcomes than FL patients. **(A)** Heatmap of TCGA HGSOC data showing unsupervised clustering of patients using the FH-and FL-specific CAF gene lists. The TCGA molecular subtypes *(28)* are shown above the heatmap in blue. Patients expressing either 75% or 50% of the FH signature genes are indicated in black and gray, respectively. Patients expressing either 75% or 50% of the FL signature genes are indicated in red and pink, respectively. **(B)** GSEA comparing the FH (left) and FL (right) gene lists to the genes expressed in TCGA mesenchymal subtype. **(C)** The ABSOLUTE (left) and ESTIMATE (right) algorithms were applied to patients falling into the FH (black), FL (red) and “other” (blue) categories of patients selected using the more stringent 75% cut-off. **(D)** Kaplan-Meier survival curves of patients falling into the FH (n=80) and FL (n=30) groups (based on the 75% cut-off). Median overall survival was 3.3 years for FH and 5.7 years for FL patients. Median disease-free survival was 1.2 years for FH and 2.1 years for FL patients. **(E)** Kaplan-Meier progression-free survival curve of patients from our centre that were profiled by flow cytometry for the proportion of CD49e+ CAFs that were positive for FAP staining. Patients were separated into FH and FL based on the median FAP expression (See Table S1B for recurrence data).

Using the ABSOLUTE algorithm, which uses somatic copy number data to estimate the cellularity of tumor samples (*29*), it was shown previously that the mesenchymal subtype in the TCGA HGSOC study had the lowest tumor purity (*30*). Using the same algorithm, FH and FL TCGA patients both had lower tumor purities than the remaining “other” patients that did not fall into either category (Figure 4C and Figure S4D). Histopathology data for the TCGA samples also indicates that tumor samples in the “other” category had a lower stromal content than both the FH and FL categories (Figure S4E). Taken together, these findings suggest that the unclassified samples could not be classified as FH or FL due to low stromal content within the tumor specimen analyzed.

ESTIMATE is another algorithm designed to estimate the quantity of infiltrating fibroblasts and immune cells using gene expression data on tumor tissues (*31*). While the ESTIMATE algorithm generated a high “stromal” score for the FH samples, it failed to identify higher stromal content in tumor samples falling into the FL category (Figure 4C and Figure S4D). Deeper analysis of this discrepancy showed that the list of 141 genes used to define “stroma” in the ESTIMATE algorithm is enriched for FH genes (21-gene overlap), but not for FL genes (2-gene overlap). As a result, patients with a high fraction of FL CAFs were not identified as having higher stromal content using this algorithm.

To determine if the CAF subtype has an impact on survival, we performed a Kaplan-Meier analysis of the TCGA patients that were classified as FH or FL (using the more stringent cut-off of 75%). FH patients had significantly shorter progression-free and overall survival than FL patients; the median overall survival of FL patients was 2 years longer than that of FH patients (**Figure 4D**). To validate this finding, we compared the %FAP+ CAFs from patients that were analyzed by flow cytometry in Figure 2F to progression-free survival for 46 of the patients for which these data were available. We defined FH patients as those with a %FAP+ CAFs above the median, and FL patients as those with a %FAP+ CAFs below the median. We found that FH patients had a significantly worse progression-free survival compared to FL patients (**Figure 4E).**

### FL cells require different culture conditions than FH cells

Most studies of CAFs utilize cultures established by plating tumor cell suspensions onto tissue culture plastic in the presence of 10% fetal bovine serum (FBS). In these conditions, CAFs rapidly adhere and proliferate, allowing for their selection and outgrowth. Our identification of CD49e as a CAF marker was facilitated by analysis of such patient-derived cultured CAF lines. Analysis of several of our cultured CAF lines indicated that they are FH by flow cytometry and express FH but not FL genes (data not shown). When we placed FH and FL cells purified by FACS directly from patient samples into standard 10% FBS conditions, we found that FH cells had a significantly higher growth rate than FL cells, and that FL cells in these conditions did not reach confluence and could not be successfully passaged (**Figure 5A)**; thus when bulk cells are placed in culture to derive CAF lines FH cells will outcompete FL cells over time due to their growth advantage. We therefore sought alternative culture conditions for FL cells; based on the expression of some genes related to adipogenesis in our FL population (e.g. *DLK1, PPARG*), we tested commercially available pre-adipocyte media and found that this media supported the expansion of FL cells over multiple passages (**Figure 5B)**. qRT-PCR analysis of FH and FL genes indicated that FL cells continued to express FL genes (*TCF21* and *DLK1*) at very high levels for up to 8 passages. However, we did see a decrease in these genes with increasing passage number, suggesting that even in pre-adipocyte media FL cells drift towards a FH phenotype with increasing time in culture (**Figure 5C**). We also saw an increase in *FAP* and *SULF1* gene expression with passage in some cases (**Figure 5C).** Thus to carry out the functional assays described below it was necessary to repeatedly isolate FL cells from patient samples and use them at passage 5 or less.

**Figure 5.**
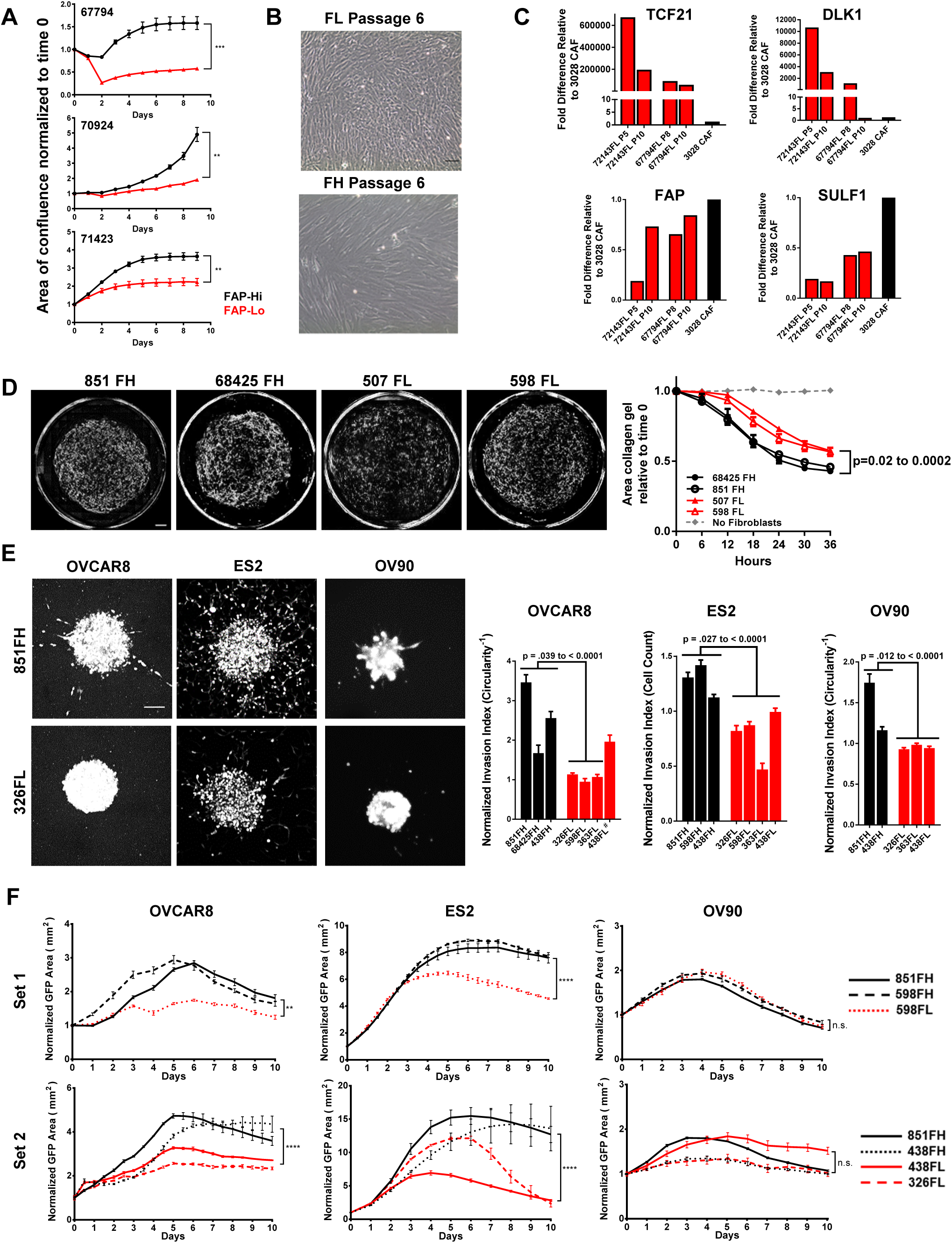
FH and FL CAFs have distinct functional properties. **(A)** Growth curves of FH and FL CAFs isolated from 3 patients cultured in 10% FBS. Mean±SEM, n=3. **p<0.01, ***p<0.001, two-way ANOVA. **(B)** FL cells grown in pre-adipocyte media. Scale bar = 50 µm. **(C)** qRT-PCR for FH genes (*FAP, SULF1*) and FL genes (*TCF21, DLK1*) in passaged FL cells (red; passage # indicated below each bar). A cultured patient-derived CAF line was used as a control (black bar). Fold-differences in gene expression were determined using the ΔΔ-Ct method, using *UBC* as a housekeeping gene, with 3 technical replicates per sample assayed. **(D)** Representative images of collagen gels cultured for 36 hours with two FH derived CAF lines and two FL-derived CAF lines (left); quantification of gel contraction over time (right). Mean ± SEM, n=4. P-values for comparisons between FH and FL lines ranged from 0.02 to 0.002, two-way ANOVA. Scale bar = 500 µm. **(E)** Representative images of spheroids generated using GFP-expressing HGSOC cell lines OVCAR8, ES2 or OV90 combined with FH or FL fibroblasts (left). Circularity was quantified OVCAR8 and OV90 cells, and the number of cells outside the spheroid was quantified for ES2 cells (right). Data is normalized to spheroids containing no CAFs. Mean±SEM, n=5 to 10 spheroids per condition. P-values calculated using Student’s t-test. Scale bar = 100 µm. #The difference between 438FL and 68425FH in OVCAR8 cells was not significant. **(F)** Growth curves of GFP-labelled cancer cells co-cultured with FH or FL CAFs after treatment with 10 µM carboplatin. Experiments were done in two separate batches (“Set 1” and “Set 2”). CAF line 851FH was used in both experiments. Data was normalized to the GFP+ area at Day 0. Mean±SEM, n=3. Asterisks reflect differences at the end of 10 days. **p<0.01, ****p<0.0001, Student’s t-test.

### FH cells promote more gel contraction, cancer cell invasion and chemotherapy resistance than FL cells

A hallmark of CAFs is their ability to contract collagen gels (*32*). Notably, two early passage FH CAF lines (851FH and 68425FH) showed a greater ability to contract collagen gels than two early passage FL CAF lines (507FL and 598FL; **Figure 5D**). Another property commonly attributed to CAFs is the ability to promote cancer cell invasion (*10, 33, 34*). To compare the ability of FH and FL CAFs to promote invasion, spheroids were established with GFP-expressing HGSOC cell lines OVCAR8, ES2 and OV90, either alone or together with FH CAFs or FL CAFs at a ratio of 5:1 (CAFs:cancer cells). The spheroids were embedded in Matrigel and imaged by fluorescence microscopy after a period of incubation at 37°C that was optimized for individual cell lines and different batches of Matrigel (**Figure S5A)**. For OVCAR8 and OV90 invasion was quantified by measuring the circularity of at least 5 spheroids per condition; a decrease in circularity is indicative of cells invading into the Matrigel, generating branches radiating away from the spheroids and thus causing spheroids to deviate from a circular shape. The invasion pattern of ES2 cells was distinct and consisted of individual cells migrating outwards into the Matrigel, rather than formation of branches. Invasion was therefore quantified by counting the number of cells outside of the sphere, rather than measuring circularity. Upon testing of FH and FL CAFs from multiple patients, FH CAFs had an overall significantly greater ability to promote the invasion of OVCAR8, ES2 and OV90 cells compared to FL CAFs. Notably, in cases where it was possible to assess patient-matched FH and FL CAFs (Patient 438 on all 3 cell lines; patient 598 on ES2 cells) there was a statistically significant difference, with FH CAFs inducing more invasion than FL CAFs (**Figure 5E**).

The standard of care for patients with HGSOC includes treatment with platinum-based chemotherapy, and chemo-resistance is a major factor leading to poor outcome in this disease. We determined the IC50s for carboplatin in OVCAR8, ES2 and OV90 cells growing under adherent conditions on plastic and showed that they ranged from 6 to 24 µM at 5 days after treatment (**Figure S6A**). Based on this information, subsequent experiments were performed using 10 µM carboplatin. To determine if FH and/or FL CAFs could influence cancer cell responses to chemotherapy, GFP-labelled OVCAR8, ES2 or OV90 cells were seeded onto feeder layers of FH CAFs or FL CAFs in flat-bottom 96 well plates. The next day 10 µM carboplatin was added and cells were cultured for an additional 10 days. GFP+ cancer cells were quantified daily using an Incucyte Zoom live cell imaging system. Both OVCAR8 and ES2 cells were rendered more resistant to carboplatin treatment in the presence of FH CAFs compared to FL CAFs, as indicated by more robust growth and a larger number of cells remaining at the end of the 10-day treatment period in FH co-cultures (**Figure 5F**). This included two pairs of patient-matched FH and FL CAFs (598FH and FL; 438FH and FL) that showed distinct outcomes. OV90 cells did not display any differences. Notably, for the two separate batches of experiments performed (**Figure 5F**) 851FH CAFs were used in both, and displayed very reproducible growth curves in all three cell lines.

### FH CAFs promote in vivo tumor growth and metastasis to lymph nodes

To ask if FH and FL CAFs have distinct influences over cancer cell behavior *in vivo*, we generated spheroids as above using luciferase-tagged OVCAR8 cells and implanted single spheroids into the mammary fat pads of non-obese diabetic/severe combined immunodeficient/IL2R-γ double-knockout (NSG) mice (**Figure S5B**). We implanted spheroids into the mammary fat pad due to earlier work by our group showing more efficient HGSOC tumor growth at this site (*22*). Tumor growth was monitored by serial imaging of luciferase activity (**Figure 6A**). Of 10 mice implanted in each group take rates were 6/10, 5/10, 6/10 and 9/10 for the no CAF control, 363FL CAFs, 374FL CAFs and 851FH CAFs, respectively, suggesting a possible enhancement of take rate in spheroids containing FH CAFs, although differences did not reach statistical significance. Tumors that contained FH CAFs (851FH) grew more rapidly compared to tumors containing FL CAFs (363FL or 374FL; **Figure 6B)**. Furthermore, 4 of 10 mice in the FH group had axillary lymph node or abdominal metastases compared to 0 of 5 mice and 1 of 6 mice in the two FL groups and 1 of 6 mice in the no CAF control group (**Figure 6C).** Importantly, this held true when the mice injected with FL CAF-containing spheroids were maintained for an additional 3 weeks, allowing the size of their tumors to reach a size exceeding the FH tumors at week 8, indicating that the difference in metastasis was not simply a function of larger tumor size in the FH group.

**Figure 6.**
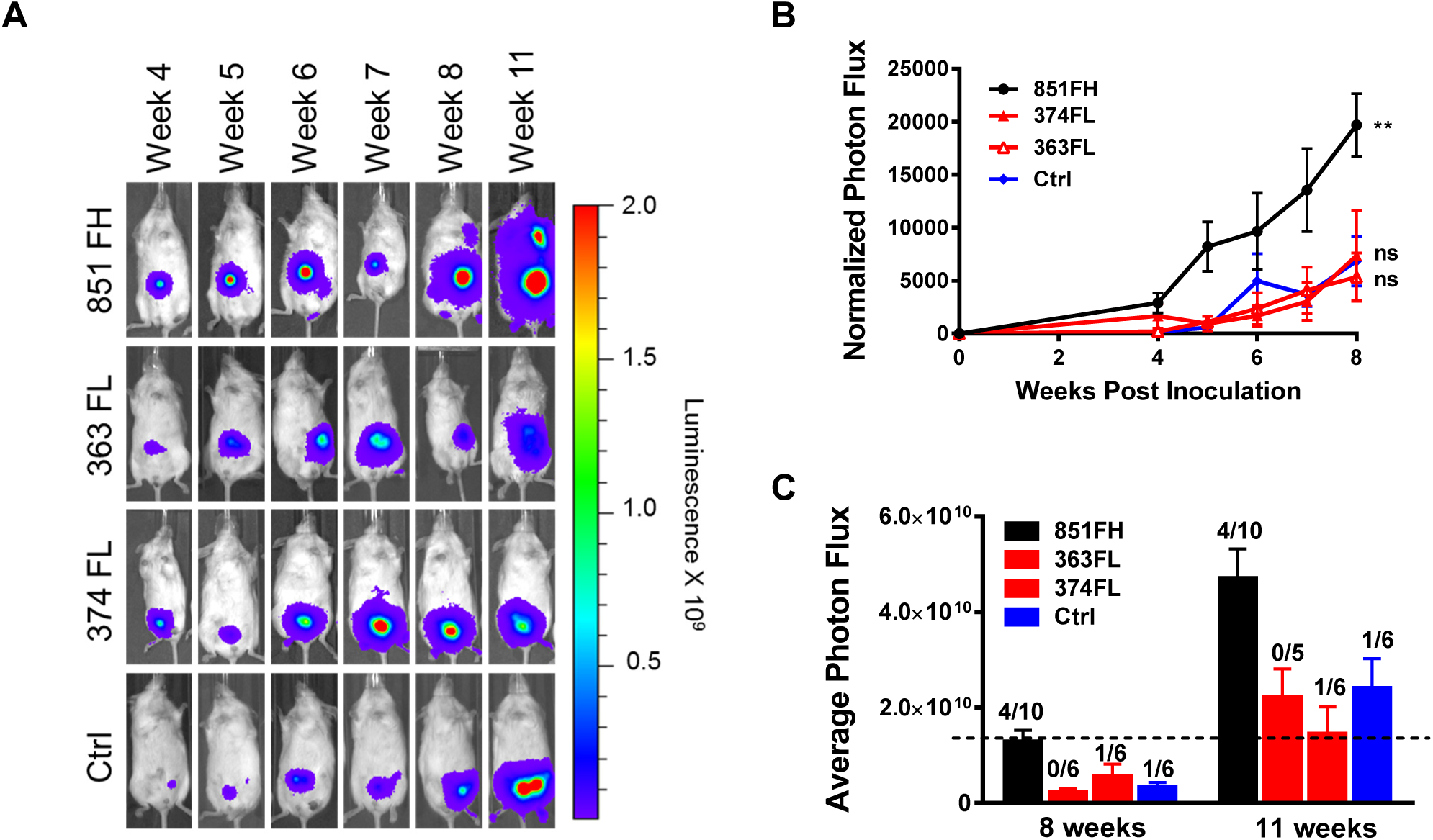
FH but not FL CAFs promote *in vivo* tumor growth and metastasis. **(A)** Spheroids of luciferase-expressing OVCAR8 cells alone or combined with FH or FL CAFs were generated as for invasion assays. After 72 hours of spheroid formation, individual spheroids were implanted into the mammary fat pads of NSG mice. Tumor growth was monitored over time using the Xenogen IVIS Imaging System 100. Representative images of a single mouse from each condition over time are shown. All mice were exposed for the same amount of time to allow visualisation of differences in tumor size over time. See also Figure S5B. **(B)** Quantification of luciferase signal, normalized to an unimplanted mouse over a period of 8 weeks. Mean±SEM, n=5 to 10 (animals in which tumors failed to grow were not included). **p<0.01, *vs* control group, linear regression. **(C)** Average tumor sizes in each group at 8 weeks and 11 weeks post-implantation. Numbers above the bars indicate the number of mice with metastases over the total number of tumor-bearing mice in each group. Mean±SEM, n=5 to 10.

### Overexpression of TCF21 in FH CAFs inhibits their pro-tumorigenic functions

TCF21 is the most highly expressed transcription factor in FL CAFs and is essential for the formation of cardiac fibroblasts during embryonic development (*35*). TCF21 has differentiation inhibiting function in skeletal muscle and smooth muscle cells (*36-38*) and is also highly expressed in white adipose tissues (*39*). Because lipid metabolism, ventricular development and cardiac related pathways were enriched in FL CAFs (**Figure 3D**), we hypothesized that TCF21 might be a master regulator of FL CAF identity. To test this possibility, 851FH CAFs were transduced with lentiviral vectors expressing either TCF21 and GFP (851FH-TCF21) or GFP alone (851FH-GFP) and GFP+ cells were FACS-purified and briefly expanded. TCF21 expression was validated by Western blot, and qRT-PCR demonstrated up-regulation of *TCF21* and two additional FL-specific transcripts (*DLK* and *TGFBR3*), as well as down-regulation of four FL-specific genes (*FAP, SULF1, MMP1* and *MFAP5*; **Figure 7A**). We then compared the functional properties of 851FH-TCF21 and 851FH-GFP cells. The ability of 851FH-TCF21 CAFs to contract collagen gels was decreased in comparison to 851FH-GFP cells (**Figure 7B).** The spheroid invasion assay was carried out using mCherry labelled OVCAR8 and OV90 cells (because the 851FH-TCF21 and 851FH-GFP CAFs were GFP+), allowing imaging of both cancer and CAF cells in this assay. The invasion of both cell types was significantly reduced upon overexpression of TCF21 in 851FH CAFs (**Figure 7C).** We next carried out co-cultures of OVCAR8, OV90 and ES2 cells with 851FH-TCF21 or 851FH-GFP CAFs in the presence of 10 µM carboplatin. OVCAR8 and ES2 cells grew more robustly in co-cultures with 851FH-GFP CAFs than in co-cultures with 851FH-TCF21 CAFs, suggesting that TCF21 expression reduced the ability of 851FH CAFs to promote the survival of cancer cells in the presence of carboplatin (**Figure 7D**). Once again, no effect was seen with OV90 cells. Finally, heterospheroids composed of OVCAR8 cancer cells and 851FH-TCF21 or 851FH-GFP fibroblasts were implanted in the mammary fat pad and growth was monitored by serial imaging of luciferase activity (**Figure 7E**). 15 of 15 mice implanted with 851FH-GFP-containing spheres grew tumors, whereas only 10 of 15 mice implanted with 851FH-TCF21-containing spheres grew tumors (p=0.042, Fisher’s exact test). In addition, overexpression of TCF21 in 851FH CAFs led to a significant growth delay compared to the 851FH-GFP control (**Figure 7F**).

**Figure 7.**
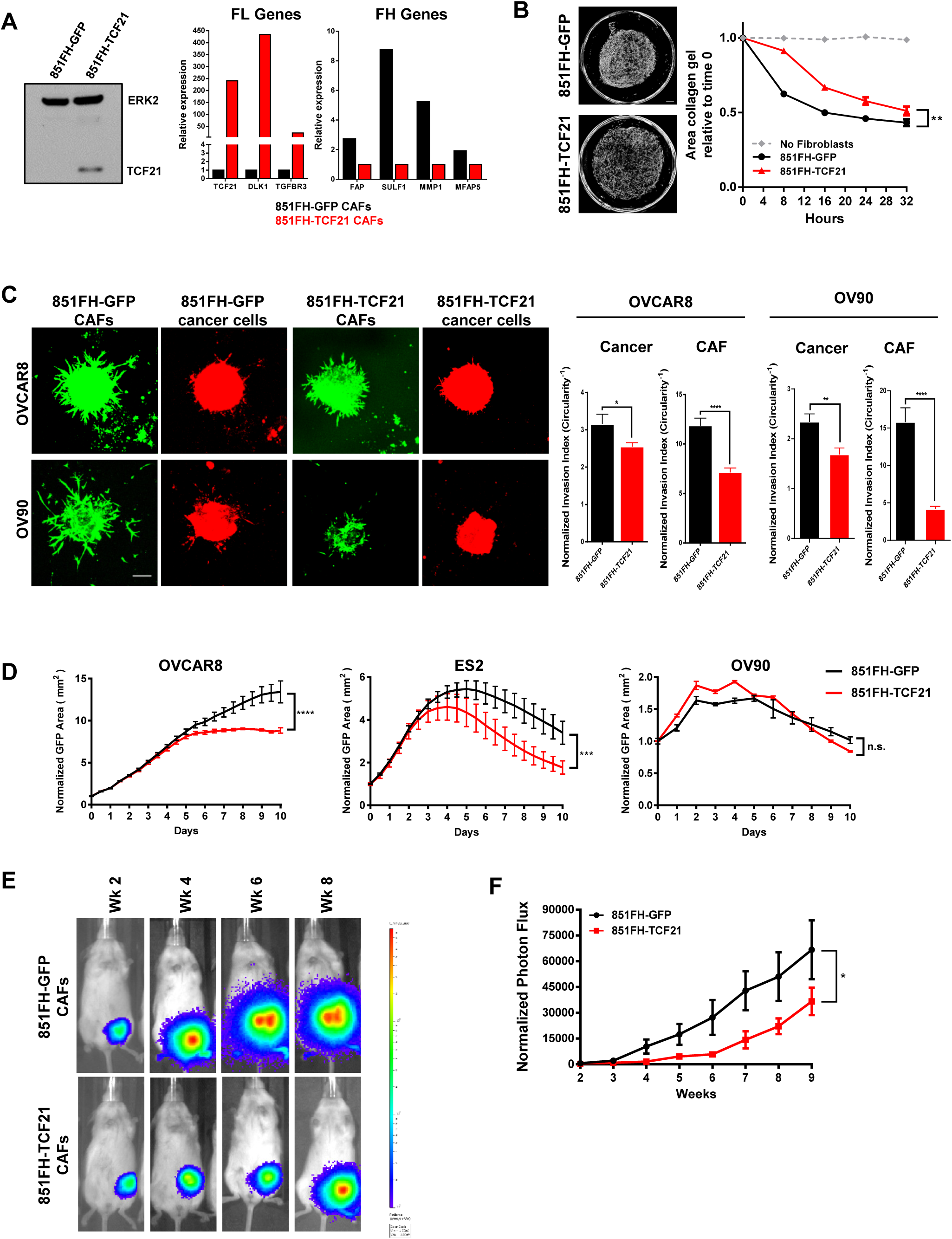
Overexpression of TCF21 in FH CAFs inhibits their pro-tumorigenic functions. **(A)** Western blot for TCF21 expression in 851FH-GFP and 851FH-TCF21 CAFs (left); and qRT-PCR for FL genes (*TCF21, DLK1* and *TGFBR3*) and FH genes (*FAP, SULF1, MMP1* and *MFAP5*) in control *vs* TCF21 overexpressing 851FH CAFs (right). **(B)** Representative images of collagen gels cultured for 32 hours with 851FH-GFP CAFs or 851-TCF21 CAFs (left); quantification of gel contraction over time (right). Mean ± SEM, n=3. **p<0.01, two-way Anova. Scale bar = 500 µm. **(C)** Representative images of spheroids generated using mCherry-expressing HGSOC cell lines OVCAR8 or OV90 with 851FH-GFP or 851FH-TCF21 CAFs mixed in. Spheroids were embedded in Matrigel and imaged after 4 days (left). Circularity was quantified in both channels (green=CAFs, red = cancer cells) using Image J (right). Mean±SEM, n=10 spheroids per condition. *p<0.05, **p<0.01, ****p<0.0001, Student’s t-test. Scale bar = 100 µm. **(D)** Growth curves of mCherry-labelled cancer cells co-cultured with 851FH-GFP or 851FH-TCF21 CAFs after treatment with 10 µM carboplatin. Data was normalized to the mCherry+ area at Day 0. Mean±SEM, n=3. Asterisks reflect differences at the end of 10 days. ***p<0.001, ****p<0.0001, Student’s t-test. **(E)** Spheroids of luciferase-expressing OVCAR8 cells alone or with 851FH-GFP or 851FH-TCF21 CAFs mixed in were generated as for invasion assays. After 72 hours of spheroid formation, individual spheroids were implanted into the mammary fat pads of NSG mice. Representative images of individual mice are shown. **(F)** Quantification of luciferase signal, normalized to an unimplanted mouse over a period of 9 weeks. Mean±SEM, n=10 to 15 (animals in which tumors failed to grow were not included). *p<0.05, two-way Anova.

## Discussion

In this study we demonstrate that CAFs in HGSOC are heterogeneous, that different subtypes have distinct influences on cancer aggressiveness and patient outcomes, and that TCF21 is a master regulator of CAF state. The identification of distinct CAF subtypes in HGSOC was facilitated by the identification of CD49e as a novel pan-fibroblast marker and the resulting ability to profile CAFs isolated directly from primary tumor specimens. We showed that patients with predominantly FH CAFs in their stroma have shorter disease-free and/or overall survival. Functional assays demonstrated that FH CAFs promote cancer cell invasion, resistance to carboplatin, and proliferation and metastasis *in vivo*, thus explaining the negative role that this CAF subtype plays in patient outcomes. By contrast, FL CAFs do not promote these behaviors in cancer cells. Finally, we show that TCF21 expression in FH CAFs suppresses their pro-tumorigenic phenotype.

While it has been known for some time that CAFs are heterogeneous (*10, 18, 40*), only recently have distinct CAF subtypes begun to be identified and characterized. For example, a recent study showed that CAFs in close proximity to cancer cells in pancreatic cancer are αSMA+ myofibroblasts, whereas fibroblasts more distant from cancer cells lack elevated αSMA and instead secrete inflammatory cytokines (*16*). A CAF subset defined by expression of CD10 and GPR77 varied in frequency in breast and lung cancers and was associated with resistance to chemotherapy and shorter patient survival (*17*). Importantly, in both of these studies the identified subtypes were all FAP-positive, suggesting that they represent heterogeneity within the FH fraction. An additional study of breast cancer used multiple fibroblast markers to identify 4 CAF subsets, of which only one was FAP-positive (“S1”) and 3 were FAP-negative (“S2-S4”) (*14*). Interestingly, the “S1” and “S4” subtypes were associated with aggressive HER2 and triple negative breast cancers, and triple negative breast cancer could be divided into two subgroups based on the presence of either S1 or S4 CAF subtypes. In a separate study the same group identified S1 and S4 CAFs in HGSOC (*15*). They showed that “mesenchymal” samples are enriched for S1 CAFs, suggesting that S1 CAFs resemble our FH CAFs. However, the S4 CAFs quickly died and could not be maintained in culture, thus the authors were unable to carry out significant characterization of these cells. In addition, no analyses were done to evaluate the clinical significance of the CAF subtypes they identified. Thus our work represents a significant advance beyond these important early studies, through demonstration of the clinical significance of CAF heterogeneity in HGSOC, as well the extensive functional and molecular characterization of isolated CAF subtypes.

The seminal studies by Givel *et al* (*15*) and Costa *et al* (*14*) showed that FAP-positive CAFs have immunosuppressive functions in HGSOC and breast cancer. Other studies have also demonstrated an immune-suppressive function of FAP-expressing stromal cells (*11-13*). We find a large cluster of pathways involved in immune regulation in FH CAFs (Figure 3D, Table S4A), which include many chemokines and cytokines involved in immune processes such as generation of myeloid-derived suppressor cells (e.g. *CXCL12, IL11, VEGF*), macrophage polarization (e.g. *CXCL12, IL10, Chi3L1*), differentiation of immune suppressive T regulatory cells (e.g. *IL1B, IL10, IL11*) and inhibition of CD8+ cytotoxic T cells (e.g. *PDCD1LG2, LGALS1*). Thus future work should include analysis of immune subtypes present in FH *vs* FL patients and understanding the cross-talk between subtypes of CAFs and immune cells in this cancer. By contrast, the FL CAFs have a distinct gene expression profile that lacks the secretory phenotype seen in FH cells. Prominent networks include translation/protein localization, ion transport/cardiac muscle contraction, and lipid metabolism and steroid biosynthesis. The latter is in agreement with our finding that FL cells grew preferentially in pre-adipocyte media, which contains low serum plus supplements that include EGF and compounds that are pro-adipogenic such as dexamethasone, 3-isobutyl-1-methylxanthine (IBMX) and ciglitazone, a PPARγ agonist. However, the prominent muscle contraction network suggests that these cells may have a more primitive mesenchymal progenitor phenotype that has both myogenic and adipogenic potential. Comparison of the FL gene signature to published mesenchymal stem cell (MSC) signatures, however, suggests that these cells are not MSCs. Indeed, MSCs more closely resemble FH CAFs as they have elevated expression of FAP as well as multiple ECM proteins and ECM remodeling enzymes (*41, 42*), thus further investigations, including functional assays, will be required to better elucidate the origin and/or identity of FL CAFs. For example, it will be of interest to compare the abilities of FH and FL CAFs to differentiate into various mesenchymal lineages and to determine if additional manipulations of the culture conditions can identify a condition that can maintain FL CAFs indefinitely *in vitro*.

Altogether, our functional assays suggest that the worse survival outcomes of patients with a FH gene signature and/or a predominance of FH CAFs within their stroma are mediated by CAF-cancer cell interactions that promote cancer cell proliferation, invasion, and therapy resistance, as well as immune-suppression as shown by others (*15*). There was some variability between individual FH CAF lines in their ability to influence these properties, and also variability in cell lines in their responses (e.g., the carboplatin response of OV90 cells was not affected by FH CAFs). This suggests different mechanisms for each of the induced behaviors, as well as different responses between individual tumors. In spite of this variation, our results indicate that future studies focused on targeting FH CAFs in order to improve outcomes and/or responses to standard chemotherapy or immunotherapy are warranted.

The unique transcriptional programs of FH and FL CAFs prompted us to more closely examine differentially expressed transcription factors between the two CAF subtypes and TCF21 was the most highly differentially expressed transcription factor in FL cells. TCF21 is expressed in epicardial progenitor cells that give rise to coronary artery smooth muscle cells and cardiac fibroblasts (*38*), the latter of which are a source of activated myofibroblasts in the infarcted heart (*43*). TCF21 is also a marker for white adipose tissue and is abundantly expressed in visceral fat-derived stem cells (*44*). When we overexpressed TCF21 in FH CAFs, our results indicated that TCF21 on its own can significantly dampen the ability FH CAFs to promote gel contraction, invasion, chemo-resistance and *in vivo* tumor growth. However, additional transcription factors or co-regulators of TCF21 and/or epigenetic regulators of chromatin accessibility are likely required to completely reprogram FH CAFs to a state that lacks pro-tumorigenic properties. Future epigenomic profiling studies will be required to compare the epigenetic states of FH and FL CAFs and identify potential avenues to “reprogram” FH CAFs to a state that is not supportive of cancer cell invasion, chemoresistance or immune suppression.

## Materials and Methods

### Primary CAF Cultures

CAFs were derived from either bulk tumor cell suspensions or FACS purified cells (see detailed methods below). Bulk tumor cell suspensions were seeded into tissue culture plates in IMDM with 10% FBS. Fibroblasts adhered and grew out preferentially under these conditions over multiple passages. Fibroblast identity was verified based on cell morphology, as well as expression of Vimentin and lack of expression of Cytokeratin. CAFs were also verified to be negative for p53 staining (i.e. wild-type) in patients with positive p53 staining (i.e. mutant) in their tumor tissues. FH CAFs isolated by flow cytometry (Viable, CD45-, CD31-, EpCAM-, CD49e+, FAP-high) were seeded into wells of 96-well culture plates in IMDM with 10% FBS. FL CAFs isolated by flow cytometery (Viable, CD45-, CD31-, EpCAM-, CD49e+, FAP-low) were seeded into wells of 96-well culture plates in pre-adipocyte media (PromoCell). Cells were passaged 1:2 when they reached confluence using 0.25% Trypsin/1 mM EDTA. All CAF experiments were done at ≤ passage 10 for cultured CAFs and FH CAFs, and ≤ passage 5 for FL CAFs.

### Cell Lines

Three HGSOC cell lines, OVCAR8, OV90 and ES2, were obtained from the ATCC and their identity was verified by Short Tandem Repeat (STR) profiling (AmpFℓSTR Identifiler, Life Technologies), performed by The Centre for Applied Genomics, The Hospital for Sick Children, Toronto, Canada, and tested negative for *Mycoplasma* infection by MycoAlert (Lonza) according to the manufacturer’s instructions. Lines were maintained as recommended by ATCC, as follows: OVCAR8 in IMDM with 10% FBS; ES2 in McCoy’s 5A media with 10% FBS; OV90 in 1:1 M199:MCDB105 media with 15% FBS.

### Mice

Female NOD/Lt-scid/IL2Rγnull (NSG) mice were bred in-house at the University Health Network Animal Resources Centre. A small incision was made below the fourth (inguinal) mammary fat pad on one. Individual spheroids suspended in 50% matrigel were drawn up into a blunt-end 18-gauge needle and implanted into the fat pad. Incisions were closed using a single wound clip.

### Primary Tumor Dissociation

Bulk tumors were mechanically minced into a slurry with sterile scalpels and then enzymatically digested in Media 199 containing 300 U/ml Collagenase and 100 U/ml Hyaluronidase mixture (Stem Cell Technologies) and 125 U/ml DNAse-I (Cedarlane) for 1-2 hours at 37°C. Following digestion, samples were centrifuged at 350 × g for 5 minutes prior to treatment with 1-2ml of ACK lysing buffer (ThermoFisher Scientific) for 5 minutes on ice. Following red cell lysis, cells were pelleted, resuspended and filtered through a 70 µm sterile nylon mesh and viable cells defined by trypan blue exclusion. Cells were cryopreserved in 90% FBS/10% dimethyl sulfoxide.

### High-Throughput Flow Cytometry (HT-FC)

HT-FC was performed on cultured CAFs and primary HGSOC tumor cell suspensions as previously described (*20*). Briefly, 363 commercially available antibodies to cell surface antigens conjugated to PE, FITC or APC were aliquoted into round bottom 96-well plates, 2 µl per well into 48 µl of flow cytometry (FC) buffer (Hanks balanced salt solution + 2% FBS). Cell suspensions of 0.5 to 1 million cells/ml in FC buffer were aliquoted by multichannel pipette into pre-prepared HT-FC plates (50 µl per well), for a final volume of 100 µl per well and a final antibody dilution of 1:50. Plates were incubated for 20 minutes on ice in the dark, centrifuged for 5 minutes at 350 x g, washed twice with 200 µl FC buffer, and resuspended in 50 to 80 µl FC buffer containing 0.1 µg/ml 4’,6-diamidino-2-phenylindole (DAPI; Sigma-Aldrich). Primary tumor samples were stained with CD45-APC-Cy7 (Biolegend, 1:200) and CD31-PE-Cy7 (Biolegend, 1:200) prior to aliquoting into plates. Fluorescence-minus-one controls were generated for each fluorochrome used and compensations were set using BD Plus CompBeads and FACSDiva software. Data collection was performed on a Becton-Dickinson LSR II flow cytometer with ultraviolet (20mW), violet (25mW), blue (20mW) and red (17mW) lasers, with default filter configuration, utilizing the High Throughput Sampler attachment. At least 10,000 events were collected per well. The gating strategy based on fluorescence-minus-one controls is illustrated in Supplementary Figure S1B.

### Flow Cytometry and FACS

Cryopreserved or fresh HGSOC single cell suspensions were washed and re-suspended at ≤1×10^7^ cell/ml in FC buffer and incubated with 10 µg/ml mouse IgG in FC buffer on ice for 10 min, followed by incubation with the following primary antibodies: CD45-PECy7 (BioLegend; 1:200), CD31-PECy7 (BioLegend; 1:200), CD49e-PE (BD Biosciences; 1:100), EpCAM-APC (BioLegend; 1:100). For experiments including CD133, an unconjugated mouse anti-human CD133 antibody was used (Miltenyi Biotec; 1:20), and for experiments including FAP, an unconjugated mouse anti-FAP antibody was used (R&D Systems; 1:50). Cells were incubated on ice for 15 minutes, washed and resuspended in FC buffer with goat-anti-mouse Alexa488 (Invitrogen; 1:400) for an additional 15 minutes, then washed and stained with the remaining directly conjugated antibodies as described above. Fluorescence-minus-one (FMO) controls were generated for each antibody and used as gating controls. Single-colour stained compensation beads (BD Biosciences) were used as compensation controls. Cells were analyzed on a BD LSR II flow cytometer or sorted using a BD FACS Aria.

### Immunofluorescence - Cytospins

FACs-sorted cells were suspended in PBS+2% FBS at a concentration of 5 × 10^4^ cells/ml and 200 µl of cell suspension were spun onto each glass slide using a cytocentrifuge at 800 rpm for 5 minutes. Slides were air-dried, fixed in 100% ice cold acetone and air dried again. Cells were permeabilized in Tris-buffered saline (TBS) containing 0.1% Tween for Cytokeratin and Vimentin, or 0.3% Triton X-100 for p53 (TBS-T) for 10 minutes, then incubated for 30 minutes in TBS-T/5% BSA/5% goat serum, followed by incubation with the following antibodies: rabbit anti-wide spectrum Cytokeratin (Abcam; 1:100), mouse-anti-human Vimentin (Abcam; 1:100), mouse-anti-human p53 (Santa Cruz; 1:100) overnight at 4°C. The following day, slides were washed 3 times in TBS and incubated with the appropriate secondary antibodies: goat-anti-rabbit-Alexa594 (Invitrogen; 1:1000) plus goat-anti-mouse-Alexa488 (Invitrogen; 1:400) for pan-CK and Vimentin; or goat-anti-mouse-Alexa594 (Invitrogen; 1:200) for p53. Secondaries were incubated for 1 hour at room temperature. Slides were then washed and incubated for 1 minute in TBS with 1 µg/ml Hoechst 333258, then coverslipped with Mowiol 4-88 (Sigma).

### Immunofluorescence - FFPE Tissue Sections

Paraffin blocks of HGSOC tissues from 5 patients were obtained from the UHN Biospecimen Sciences Program (UHN, Toronto, ON) in accordance with regulations for excess tissue use stipulated by the UHN research ethics board. 4 µm sections were transferred onto positively-charged slides. Sections were deparaffinised using xylene and ethanol. For antigen retrieval, slides were incubated in 0.01 M citrate buffer (pH 6.0) with 0.05% Tween in a glass vessel submerged in boiling water for 20 minutes. Sections were then permeabilized in TBS-T (containing 0.1% Tween for Cytokeratin, CD49e and FAP, and 0.3% Triton X-100 for TCF21) for 10 minutes. Sections were incubated for 2 hours in TBS-T/0.5% BSA/5% goat serum, followed by incubation with rabbit anti-human TCF21 (Sigma, 1:100), mouse-anti-human-CD49e (BD Biosciences, 1:100), mouse-anti-human-FAP (R&D Systems, 1:50), and either mouse-anti-pan-Cytokeratin (Abcam, 1:100), or rabbit anti-wide spectrum Cytokeratin (1:100) overnight at 4°C. The following day, slides were washed 3 times and incubated with goat-anti-rabbit-Alexa594 (Invitrogen; 1:1000) plus goat-anti-mouse-Alexa488 (Invitrogen; 1:400) for 1 hour at room temperature. Slides were then washed and incubated for 1 minute in TBS with 1 µg/ml Hoechst 333258, then coverslipped with Mowiol 4-88 (Sigma). Slides were scanned using a Zeiss Axio Slide Scanner and images were analyzed using HALO software (Indica Labs).

### Microarrays

HGSOC single cell suspensions were stained for fluorescence activated cell sorting (FACS) as described above. Doublets and dead cells were excluded and CD31^-^CD45^-^EpCAM^-^ CD49e^+^ cells, CD31^-^CD45^-^EpCAM^+^CD133^-^ cells, and CD31^-^CD45^-^EpCAM^+^CD133^+^ cells were gated for sorting based on FMO controls. Post-sort purity checks for each sample confirmed >98% purity for each population. Twelve patient samples were sorted into tubes containing Iscove’s Modified Dulbecco’s Medium (IMDM) with 10% FBS. Sorted populations were washed with PBS and then RNA was extracted immediately using the RNeasy Plus Micro kit (Qiagen). RNA quality was verified using a Bioanalyzer 2100 (Agilent Technologies). All samples had an RNA integrity number (RIN) >8. 5 ng of RNA per sample were amplified using the Ovation pica WTA V2 kit (Nugen) as per the manufacturer’s instructions. Amplified cDNA from each sample was labelled following Nugen Illumina solution application Note #2. 750 ng of amplified biotin-labelled cDNA generated from these samples were randomized and hybridized onto three Illumina Human HT-12 v4 BeadChips. BeadChips were incubated at 48 °C at rotation speed 5 for 15 hours for hybridization. The BeadChips were washed and stained as per Illumina protocol and scanned on the iScan (Illumina). Data files were quantified in GenomeStudio Version 2011.1 (Illumina) and passed sampled-dependent and independent QC metrics.

Probe intensities were normalized between arrays using quantile normalization and transformed using the logarithm of base 2. Differential expression between the different groups of samples – CD49e+ and EpCAM+, FH and FL – was estimated using limma 3.28.21. HGNC gene names were associated with probe identities using biomaRt 2.28.0. Two-color heatmaps were generated using the heatmap.2 function of the gplots R package 3.0.1 and pvclust 2.0.0 was used to cluster the CAF FH and FL samples with bootstrapping.

To carry out pathway analysis of CD49e+ and EpCAM+ cells, an expression score was created to rank all genes from top up-regulated to top down-regulated using the formula ‘sign(logFC) * -log10(pvalue)’. Gene set enrichment analysis (GSEA – Broad Institute) was applied on this rank list using 1000 permutations. The gene-sets tested by GSEA were the Reactome database included in the MSig c2.cp.reactome.v6.1.symbols.gmt file. Results were visualized in Cytoscape 3.6.1 using EnrichmentMap 3.1, clustered and annotated using AutoAnnotate 1.2. Cluster labels were manually edited for clarity.

### qRT-PCR

Quantitative reverse transcription PCR (qRT-PCR) was performed in triplicate using Power SYBR Green PCR Master Mix (Life Technologies). Samples were loaded into a BioRad CFX96 real-time PCR detection system following the manufacturer’s protocols. Relative amounts of mRNA were calculated by the ΔΔCt method and normalized to expression levels of *UBC.* The following primer sequences were used: *FAP*, forward 5′-TGGCGATGAACAATATCCTAGA-3′, reverse 5′-ATCCGAACAACGGGATTCTT-3′; *Col11a1*, forward 5′-TTTTCCAGGATTCAAAGGTGA-3′, reverse 5′-TGGGCCAATTTGACCAAC-3′; *Sulf1*, forward 5′-ACCAGACAGCCTGTGAACAA-3′, reverse 5′-ATTCGAAGCTTGCCAGATGT-3′; *TCF21*, forward 5′-CGACAAATACGAGAACGGGTA−3′, reverse 5′-TCAGGTCACTCTCGGGTTTC−3′; *Colec11*, forward 5′-CCCCTGGTCCTAATGGAGA−3′, reverse 5′-TCAGCTGAGAGACCTGGTTGT−3′; *Dlk1*, forward 5′-GACGGGGAGCTCTGTGATAG−3′, reverse 5′-GGGCACAGGAGCATTCATA−3′; *UBC*, forward 5′-AGGCAAAGATCCAAGATAAGGA-3′, reverse 5′-GGACCAAGTGCAGAGTGGAC−3′.

### RNA-Seq

HGSOC single cell suspensions were stained for FACS as described above. Doublets and dead cells were excluded and CD45-CD31-EpCAM-CD49e+FAP-High and CD45-CD31-EpCAM-CD49e+FAP-Low populations were gated for sorting. At least 10000 cells were sorted from each population. Twelve patient samples were sorted into tubes containing Iscove’s Modified Dulbecco’s Medium (IMDM) with 10% FBS. Sorted populations were washed with PBS and then RNA was extracted immediately using the RNeasy Plus Micro kit (Qiagen). RNA samples were assessed on a RNA 6000 Pico chip (Agilent Technologies) using the Agilent Bioanalyzer to determine sample RIN and quantified by the Qubit RNA HS assay kit (Life Technologies). All samples used had RIN values greater than 8.5. 4 ng of RNA were used to prepare RNA libraries using the SMARTer Stranded Total RNA-seq Kit – Pico Input Mammalian (Takara Bio USA Inc.). Briefly, the samples underwent first-strand synthesis *via* random priming oligos on the 3’ end. A template switching oligo mix (TSO) was then incorporated to allow the RT reaction to continue replicating the 5’ of the RNA strand. Following this, the samples were PCR amplified to incorporate full-length Illumina adapters and sample barcodes by binding to either the TSO stretch on the 5’ end, or the random priming oligo sequence on the 3’ end. The amplified cDNA is treated with ZapR which specifically targets ribosomal RNA in the presence of mammalian-specific R-Probes. This process leaves non-ribosomal RNA untouched, while ribosomal RNAs are cleaved, leaving them non-amplifiable. A final PCR reaction was done to enrich the uncut strands of cDNA to generate the final RNA library. Final RNA library sizing was verified on the Agilent high sensitivity DNA kit (Agilent Technologies) using the Agilent Bioanalyzer while library concentration was quantified by qPCR using the KAPA SYBR FAST qPCR kit (Kapa Biosystems). Libraries were normalized to 10nM, diluted to 2nM, denatured using 0.2N NaOH, and diluted again to 1.7pM before loading onto the NextSeq 500 system. The samples were sequenced using a paired-end 75 cycle sequencing run to achieve a minimum of ∼40M reads per sample.The reads were mapped using STAR/2.5.2 to the hg38 reference genome. Read counts per gene were obtained through htseq-count v.0.6.1. After removing low count genes whose cpm (counts per million reads) were less than 0.75 to in one third of the total number of samples, the edgeR R package v.3.8.6 was used to normalize the data using the TMM (trimmed mean of M values) method and to estimate differential expression by applying a generalized linear model between the FL fraction samples and the FH fraction samples. The multidimensional scaling plot was created using the edgeR plotMDS.DGEList() function.

For pathway analysis of FH and FL CAFs, an expression score was created to rank all genes from top up-regulated to top down-regulated using the formula sign(logFC) * - log10(pvalue). Gene set enrichment analysis (GSEA – Broad Institute) was applied on this rank list using 1000 permutations. The gene-sets tested by GSEA were first the microarray FH and FL gene lists and second gene-sets from the Reactome database included in the MSig c2.cp.reactome.v6.1.symbols.gmt file. Results were visualized in Cytoscape 3.6.1 using EnrichmentMap 3.1, clustered and annotated using AutoAnnotate 1.2. Cluster labels were manually edited for clarity.

### TCGA Data Analysis

To identify genes specific to FH and FL CAFs, genes differentially expressed between FH and FL fractions at a FDR cut-off of 0.05 and with a logFC greater than 2 fold (logFC >2 for FH and <-2 for FL) were selected. These genes were then filtered to include only those that have a higher expression in CD49e+ cells compared to EpCAM+ CD133+ at a FDR cut-off of 0.05 in the Human Illumina HT-12 V4 array. This generated a list of 165 FH CAF-specific genes and 78 FL-specific genes.

HGSOC data were downloaded from the Genomic Data Commons (GDC) Data Portal (https://portal.gdc.cancer.gov/). HiSeq gene level counts (level2 RNA-Seq data) and corresponding clinical data were downloaded for 374 samples included in this analysis.

Trimmed mean of M values (TMM) followed by count per million (CPM) and logarithm of base 2 transformation was used to normalize the data within the edgeR package. Data were selected to only contain the FH and FL gene list and a heatmap was created using R heatmap.2. Color bars were added to the heatmap to identify patients based on molecular phenotype described by Verhaak *et al* (*28*). These phenotypes include the categories ‘mesenchymal’, ‘immune’, ‘proliferative’, and ‘differentiated’ and they were extracted from Supplemental Table 1 of Verhaak *et al* (*28*). Differential expression was calculated within edgeR for samples defined as ‘mesenchymal’ versus all the other samples and a list ranking genes from top up-regulated to down regulated was generated using the formula ‘sign(logGC) * -log10(pvalue)’.

FH and FL color bar: TCGA patients were ranked using a score that counts how many genes from the FH gene list or FL gene list have a normalized value greater than the patient mean (i.e. a z-score). Patients were considered FH if they had positive scores for at least 75% of the gene list (corresponding to a sum of z-scores > gene list length/2). A less tringent threshold classified patients as FH or FL if they had positive scores in at least 50% of the gene list (corresponding to a sum of z-scores > 0). The FH and FL patient categories were added to the heatmap color bar and used for further analysis.

TCGA patients were then grouped into ‘FH’ and ‘FL’ and ‘other’ categories using either the 75% or the 50% thresholds and ESTIMATE and ABSOLUTE values, which are known to measure percentage of stromal cell and tumor purity content, respectively (*29, 31*) were determined using the respective R packages. Data were plotted on a whisker boxplot for each category. Clinical data including overall survival and progression free events were retrieved for TCGA patients falling into the FH and FL categories based on the more stringent 75% cut-off and Kaplan-Meier curves were generated and associated with a log rank test using GraphPad Prism software.

For GSEA testing of the short FH and FL lists in the ‘mesenchymal’ subtype, the mesenchymal rank list was tested against the short FH and FL gene lists using default parameters.

### Gel Contraction

FH and FL CAFs were suspended in IMDM with 10% FBS at a concentration of 100,000 cells/ml and kept on ice. 3 mg/ml Collagen-I solution (Thermo-Fisher Corning, 354236) and 1M NaOH were then added to the cell suspension give a final concentration of 1mg/ml Collagen-I and a neutral pH, and 100 µl were aliquotted into non-tissue culture-treated flat-bottom 96-well plates. Plates were left for 20 minutes at room temperature to solidify, then 120 µl IMDM + 10% FBS was added to the wells. A small pipet tip was gently run around the perimeter of each well prior to imaging on the Incucyte ZOOM every 6 hours for 3 days. The areas of the gels at different time points were quantified using ImageJ (Fiji) software.

### 3D Spheroid Invasion Assays

GFP-expressing HGSOC cell lines (OVCAR8, OV90 and ES2) were mixed with either FH, FL, 851FH-TCF21, or 851FH-GFP CAFs at a ratio of 5:1 (CAF: cancer cells). 6000 total cells per well were then plated in 90 µl basal media (IMDM for OVCAR8, McCoy’s 5A for ES2, or MCDB105: M199 for OV90) supplemented with 2% FBS in ultra-low attachment round bottom 96 well plates (Corning, 7007). Plates were centrifuged at low speed to center the cell suspension prior to incubation at 37°C for 3 days. At that point, spheroids were formed and 30 µl of media were removed from each well and replaced with 30 µl of growth factor reduced Matrigel (Corning, 354230) to give a final Matrigel concentration of 33% v/v. Plates were incubated for 60 minutes at 37°C, then 100 µl per well of the appropriate basal media supplemented with 2% FBS was added and plates were returned to the incubator. The time point used to quantify invasion was optimized for individual cell lines, and ranged from 1 day (for ES2 cells) to 4 days (for OVCAR8 and OV90 cells). Spheroids were imaged with a Zeiss LSM700 confocal microscope. Circularity of spheroids (OVCAR 8, OV90) and cell counts outside the spheroid (ES2) were analyzed using ImageJ (Fiji) software. In some cases, multiple spheroids formed per well, or the spheroids were not centered in the well and thus could not be imaged. However, a minimum of 5 and up to 10 spheroids per experimental group were analyzed in all cases.

### In Vitro Carboplatin Treatment

1000 cells per well of GFP or mCherry-expressing OVCAR8, OV90 or ES2 cells were seeded in their respective media onto a monolayer of CAFs in 96-well flat-bottom tissue culture plates. The plates were incubated at 37°C for 24 hours, then carboplatin (Hospira) was added to reach a final concentration of 10 µM. Cancer cell proliferation was monitored using Incucyte ZOOM live cell imaging system for up to 10 days.

### In Vivo Assays

HGSOC OVCAR8 cells constitutively expressing firefly luciferase were mixed with FH, FL, 851FH-TCF21 or 851FH-GFP CAFs at a ratio of 5:1 (CAF: cancer cells). OVCAR8 cells alone were used as a control. Spheroids were formed as described above and suspended in a final volume of 100 µl IMDM media supplemented with 2% FBS. 100 µl of growth factor reduced Matrigel was added to the wells, mixed gently, then immediately loaded into blunt-end 16 gauge syringes (Stemcell Technologies, 28110). 6-8 week old female NSG mice were anaesthetized with isoflurane and an incision was made near the fourth mammary fat pad. Spheroids were then implanted into the fat pad directly and the incision was stapled. Mice were injected weekly with 30 mg/ml luciferin (Cedarlane) and imaged using a Xenogen IVIS Imaging System 100. Signal was quantified using IVIS Living Image software.

### Lentiviral Constructs

Custom lentiviral vectors were designed and plasmids obtained using VectorBuilder (Cyagen Biosciences). A PGK promoter-driven *TCF21*-P2A-GFP lentiviral vector and control PGK-GFP and PGK-mCherry vectors were designed and purchased from Vectorbuilder (Cyagen Biosciences). For lentiviral infections cells were plated in 6-well plates at 1.0×10^5^ cells per well and incubated with viral supernatants for 48 hours at 37°C. Infected cells were purified by FACS on the basis of GFP or mCherry expression and expanded for further use.

### TCF21 Western Blot

851FH-GFP and 851FH-TCF21 CAFs were lysed in RIPA lysis buffer supplemented with EDTA-free protease inhibitor cocktail tablets (Roche) and normalized for total protein amount. 35 µg of protein from each sample were resolved in a 12% SDS-PAGE gel, and transferred onto Immobilon-P membranes (Millipore) using a semi-dry transfer method (Bio-Rad). Blots were probed overnight at 4°C using a mouse-anti-human ERK2 antibody (Santa Cruz; 1:1000) and a rabbit-anti-human TCF21 antibody (Sigma; 1:250), followed by a 45-minute incubation at room temperature with a goat anti-mouse IgG HRP-linked secondary antibody (Invitrogen, 1:1000), and goat-anti-rabbit IgG HRP-linked secondary antibody (Cell Signaling, 1:2500). Proteins were detected using enhanced chemiluminescence reagent (ThermoFisher) and autoradiograph exposure (Sigma-Aldrich).

### Quantification and Statistical Analysis

Information about statistical details and analysis of microarray and RNAseq data is indicated in text, figure legends, or method details. Graphs and statistical values (p-values, correlation coefficients, hazard ratios) were generated using Graphpad Prism 6.03. Error bars indicate standard error of the mean (SEM) or standard deviation (SD) for a minimum of three independent experiments.

### Data Availability

Microarray and RNA-seq data that support the findings of this study have been deposited at NCBI’s Gene Expression Omnibus and are accessible through GEO Series accession number GSE126133.

### Study Approval

Tumor samples were obtained from 69 patients with high grade serous ovarian cancer who underwent surgery at the University Health Network (**Table S1a**). All patient tumor samples were collected after obtaining written informed consent according to the research protocol #06-0903, approved by the University Health Network Research Ethics Board, Toronto, Canada. Animal experiments were performed in accordance with national and institutional guidelines approved by the Canadian Counsel on Animal Care and approved by the University Health Network Animal Care Committee protocol #1542.

## Supplementary Materials

Figure S1. Identification of CD49e as a CAF-specific cell surface marker

Figure S2. Isolation and transcriptional profiling of CD49e+ CAFs

Figure S3. IF staining and quantification of FAP and TCF21 in HGSOC specimens

FigureS4. FH and FL gene lists and FH/FL TCGA patient classification

Figure S5. Schematics for *in vitro* and *in vivo* spheroid assays

Figure S6. Chemotherapy treatment of HGSOC cell lines

Table S1A. Clinical Data and %FAP+ CAFs in Primary Tumor Specimens

Table S1B. Recurrence Data for Patients in Figure 4E.

Table S2A - Differentially Expressed Genes Between FACS Isolated CD49e+ CAFs and EpCAM+CD133+ Epithelial Cells

Table S2B - Differentially Expressed Gene Sets (MsigDB Reactome) Between FACS Isolated CD49e+ CAFs and EpCAM+CD133+ Epithelial Cells

Table S3A - Differentially Expressed Genes Between FH and FL Clusters of Isolated CD49e+ CAFs

Table S3B – Differentially Expressed Genes Between FL CAFs and Normal Stroma in the Leung et al Dataset.

Table S4A – Differentially Expressed Genes Between FACS-Purified FH and FL CAFs

Table S4B - Differentially Expressed Gene Sets (MsigDB REactome + GO BP and MF) Between FACS-Purified FH and FL CAFs

Table S5 – Gene Signature Used to Interrogate TCGA Data

## Supporting information

Table S1A - Clinical data all patients

Table S1B - Recurrence data for patients in Figure 4E

Table S2A - CD49e vs EpCAM gene list

Table S2B - CD49e vs EpCAM GSEA results

Table S3A - FH vs FL geneslist microarray

Table S3B - FH vs normal gene list from Leung et al study

Table S4A - FH vs FL genelist RNAseq

Table S4B - FH vs FL GSEA results

Table S5 - FH FL gene signature

## Acknowledgements

We thank the patients who donated their cancer tissues for research, and the UHN Program for Biospecimen Sciences for facilitating fresh tissue acquisition. We are also grateful to the UHN Animal Resources Centre staff for animal care. Thanks also to Catherine O’Brien and Evelyne Lima Fernandes for critical reading of the manuscript.

## Author contributions

AH, BGN and LEA designed research studies, AH, SP, JM, JD and JP conducted experiments and acquired data, VV, GDB and MQB analyzed data, VWH, KHT, BC and MQB provided reagents, including clinical specimens; AH, BGN and LEA wrote the manuscript.

