## Supplemental figures for "Distinct cancer-associated fibroblast states drive clinical outcomes in high-grade serous ovarian cancer and are regulated by TCF21"

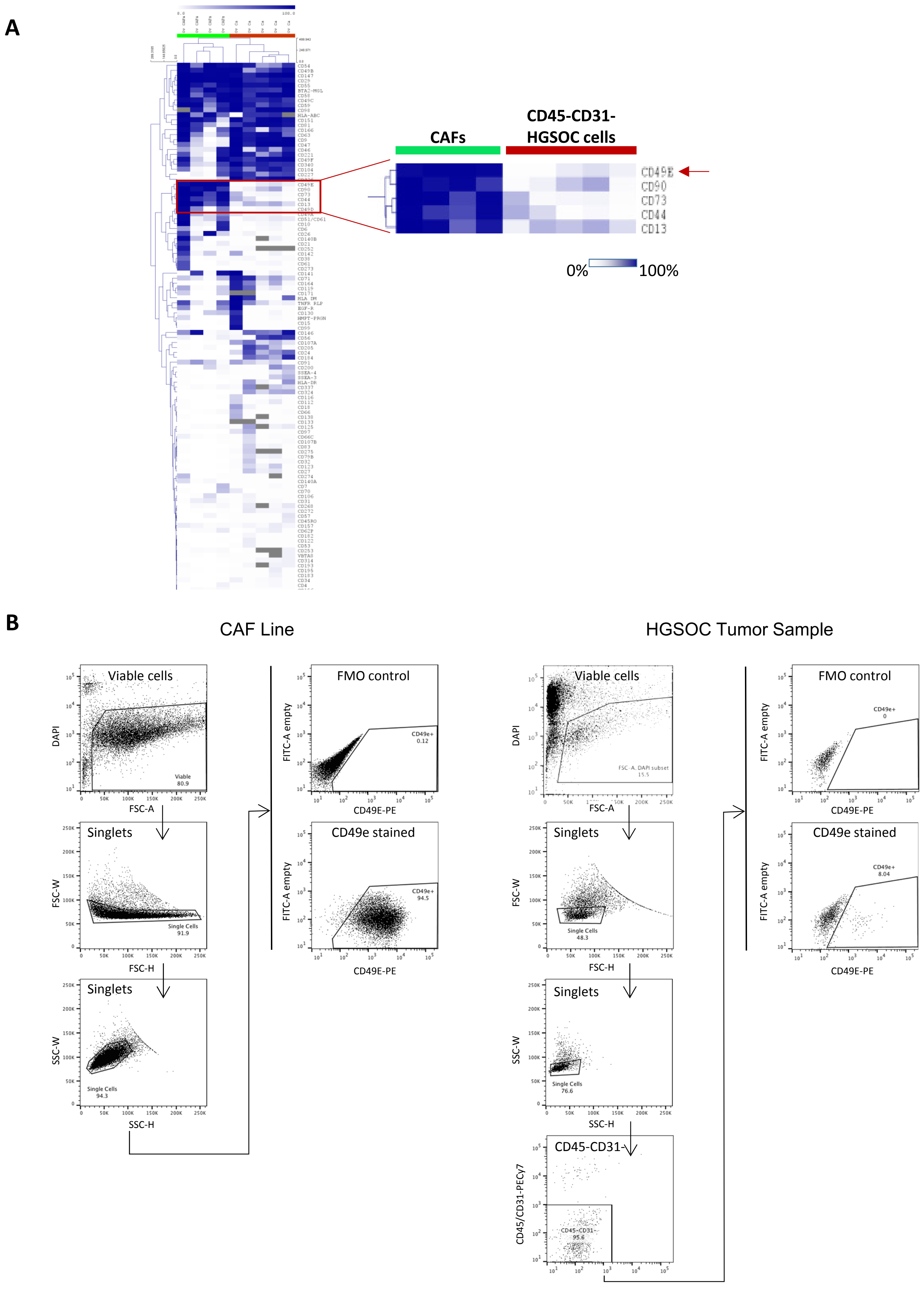
A
CAFs
CD45-CD31-
HGSOC cells
CD49E
CD90
CD73
CD44
CD13
0%
100%
B
CAF Line
HGSOC Tumor Sample
Viable cells
Singlets
Singlets
FMO control
CD49e stained
CD45-CD31-
DAPI
FSC-A
FSC-W
FSC-H
SSC-W
SSC-H
FITC-A empty
CD49E-PE
CD45/CD31-PECy7

**Figure S1. Identification of CD49e as a CAF-specific cell surface marker. (A)** We performed high throughput flow cytometry with panel of 363 antibodies on 4 CAF lines and 5 HGSOC patient samples. Patient samples were co-stained with CD45 and CD31 to exclude inflammatory and endothelial cells from the analysis. A heat map of percent-positive cells for each antibody on CAFs and the CD45-CD31- fraction of tumor samples was generated using unsupervised hierarchical clustering with a Pearson correlation distance metric and complete linkage. A cluster of differentially expressed markers was found, of which CD49e was the top hit. **(B)** Representative FACS plots of CAFs (left) and a primary HGSOC sample (right) stained for CD49e. The gate for CD49e expression was set based on a fluorescence-minus-one (FMO) control. Related to Figure 1.

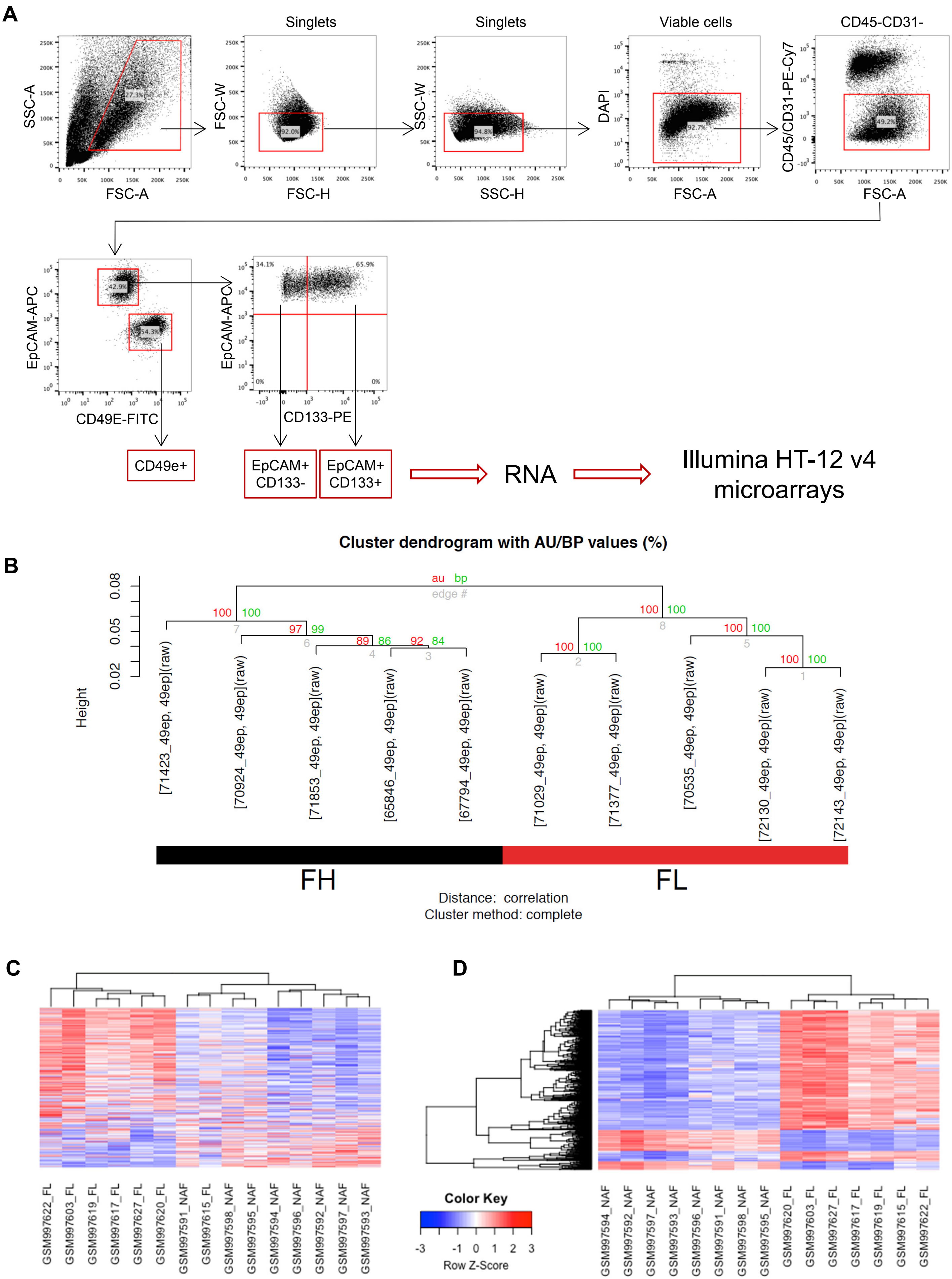
A
Singlets
Singlets
Viable cells
CD45-CD31-
SSC-A
FSC-A
FSC-W
FSC-H
SSC-W
SSC-H
DAPI
FSC-A
CD45/CD31-PE-Cy7
FSC-A
27.3%
92.0%
94.8%
92.7%
49.2%
EpCAM-APC
CD49E-FITC
42.9%
54.3%
EpCAM-APC
CD133-PE
34.1%
65.9%
0%
0%
CD49e+
EpCAM+ CD133-
EpCAM+ CD133+
RNA
Illumina HT-12 v4 microarrays
Cluster dendrogram with AU/BP values (%)
B
au bp
edge #
Height
[71423_49ep, 49ep](raw)
[70924_49ep, 49ep](raw)
[71853_49ep, 49ep](raw)
[65846_49ep, 49ep](raw)
[67794_49ep, 49ep](raw)
[71029_49ep, 49ep](raw)
[71377_49ep, 49ep](raw)
[70535_49ep, 49ep](raw)
[72130_49ep, 49ep](raw)
[72143_49ep, 49ep](raw)
FH
FL
Distance: correlation
Cluster method: complete
C
GSM997622_FL
GSM997603_FL
GSM997619_FL
GSM997617_FL
GSM997627_FL
GSM997620_FL
GSM997591_NAF
GSM997615_FL
GSM997598_NAF
GSM997595_NAF
GSM997594_NAF
GSM997596_NAF
GSM997592_NAF
GSM997597_NAF
GSM997593_NAF
D
Color Key
Row Z-Score
GSM997594_NAF
GSM997592_NAF
GSM997597_NAF
GSM997593_NAF
GSM997596_NAF
GSM997591_NAF
GSM997598_NAF
GSM997595_NAF
GSM997620_FL
GSM997603_FL
GSM997627_FL
GSM997617_FL
GSM997619_FL
GSM997615_FL
GSM997622_FL

**Figure S2. Isolation and transcriptional profiling of CD49e+ CAFs. (A)** Full gating strategy for isolation of cell populations used in microarray analysis. **(B)** Cluster dendrogram of the isolated CD49e+ CAFs isolated from 10 HGSOC samples (Human Illumina V4 array) after bootstrapping. Red and green numbers are the result of the pvclust function to assess the uncertainty in hierarchical cluster analysis. Red font corresponds to the AU (approximately unbiased) p-value and green font to the BP (bootstrap probability) value for each cluster in the dendrogram. 100 (100%) means that the same cluster was obtained for all bootstrapping permutations. Related to Figure 1E. **(C)** Heatmap of the top 500 FL genes (same as in Figure 2A) showing only the FL and normal stroma samples from the Leung *et al* study, demonstrating that the majority of the FL genes are higher in FL stroma *vs* normal adjacent stroma. **(D)** Heatmap of the top 500 probes from the comparison of the Leung *et al* FL stroma *vs* normal adjacent stroma samples. Related to Figures 1 and 2.

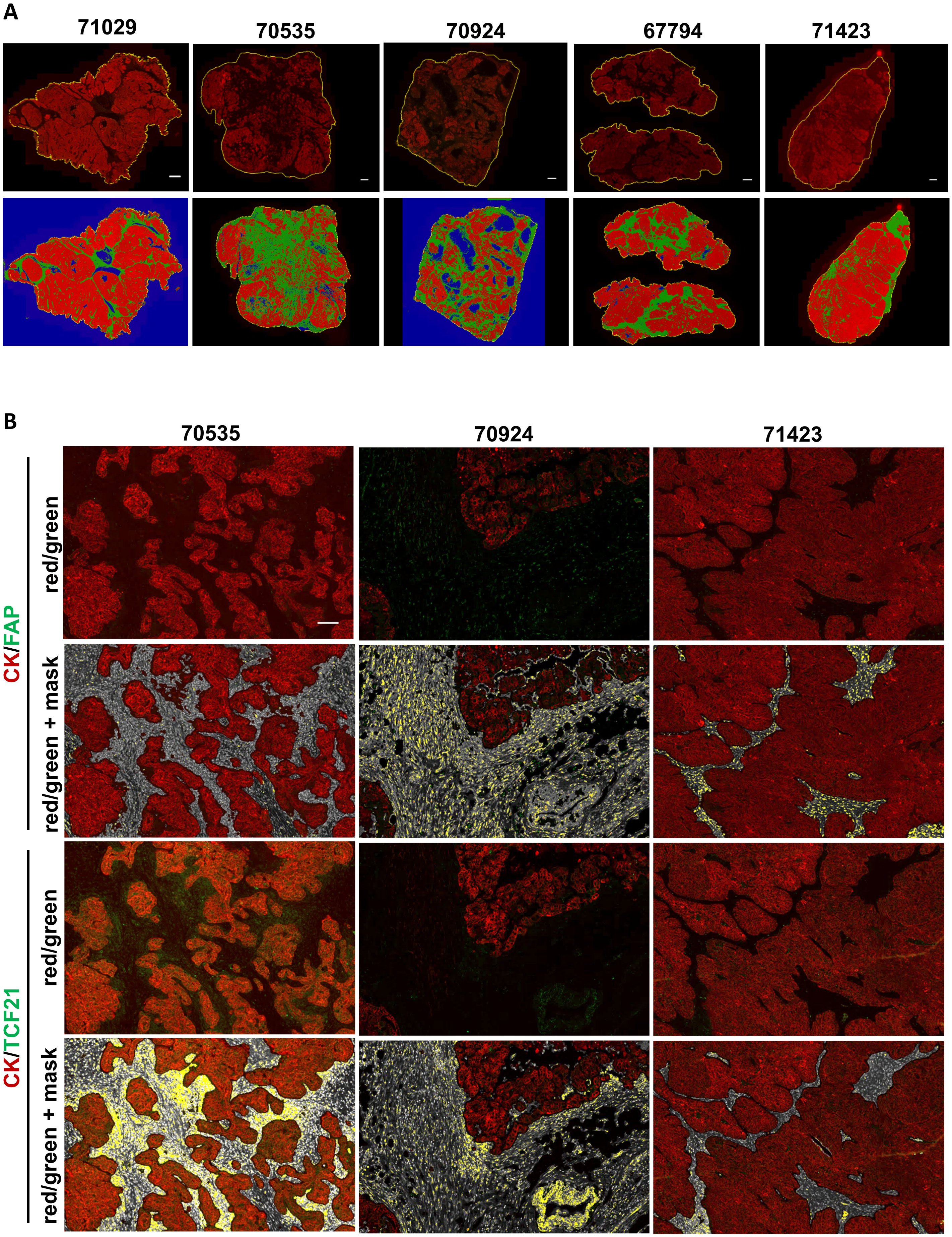

A
71029
70535
70924
67794
71423
B
70535
70924
71423
CK/FAP
red/green
red/green + mask
CK/TCF21
red/green
red/green + mask

**Figure S3. IF staining and quantification of FAP and TCF21 in HGSOC specimens. (A)** Low power images of entire tissue sections showing the pan-CK staining in the red channel (top row) and the HALO software-generated classification of epithelial (red) and stromal (green) compartments. Blue regions are empty space. Scale bar = 1 mm. **(B)** Representative images of HGSOC specimens at higher magnification showing the epithelial pan-CK-positive (red) and FAP- or TCF21-positive (green) staining and the corresponding mask generated by HALO software for quantification of the FAP or TCF21 signal within the stromal regions. Serial sections are shown for FAP *vs* TCF21 comparison. DAPI-positive nuclei within the stroma are white, and FAP or TCF21 positive signal in the green channel are overlayed with yellow. Scale bar = 500 μm. Related to Figure 2C-E.

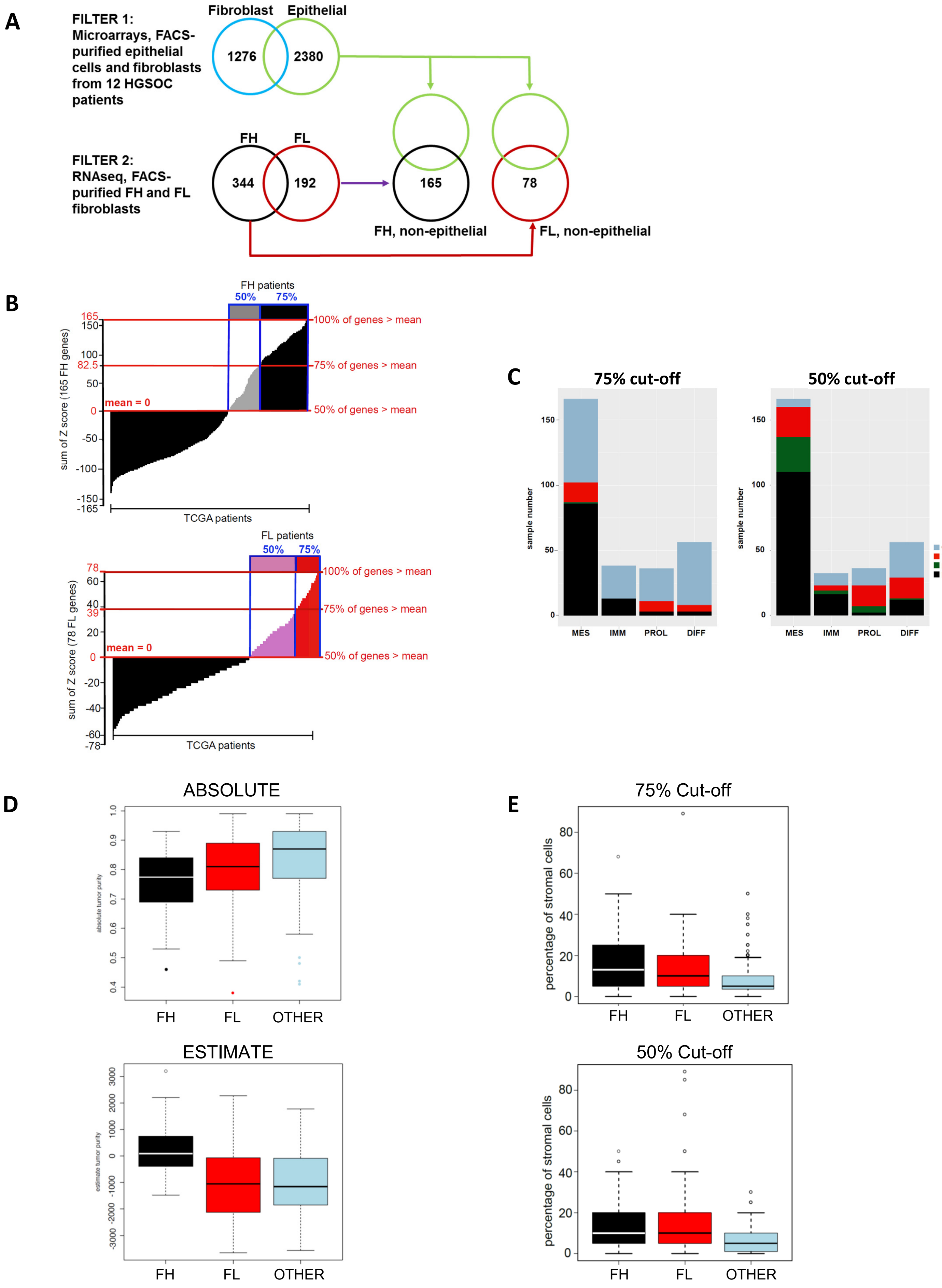
A
FILTER 1:
Microarrays, FACS-purified epithelial cells and fibroblasts from 12 HGSOC patients
Fibroblast
Epithelial
1276
2380
FILTER 2:
RNAseq, FACS-purified FH and FL fibroblasts
FH
FL
344
192
165
78
FH, non-epithelial
FL, non-epithelial
B
FH patients
50%
75%
100% of genes > mean
75% of genes > mean
50% of genes > mean
mean = 0
sum of Z score (165 FH genes)
TCGA patients
FL patients
sum of Z score (78 FL genes)
C
75% cut-off
50% cut-off
sample number
MES
IMM
PROL
DIFF
other
FL
FH&FL
FH
D
ABSOLUTE
absolute tumor purity
FH
FL
OTHER
ESTIMATE
estimate tumor purity
E
75% Cut-off
50% Cut-off
percentage of stromal cells

**Figure S4. FH and FL gene lists and FH/FL TCGA patient classification**. **(A)** Schematic of filtering strategy used to generate a gene signature for interrogation of TCGA data. Filter 1 is selecting genes that are differentially expressed in CD49e+ cells compared to cancer epithelial cells (FDR ≤0.05) and filter 2 is selecting genes based on the comparison of purified FH and FL CAFs (FDR ≤0.05 and logFC ≥2). **(B)** TCGA patients were ranked using a score that counts how many genes from the FH gene list (top) or FL gene list (bottom) have a normalized value greater than the patient mean (*i.e.* a z-score). Patients with positive scores in at least 75% of the gene list (corresponding to a sum of z-scores ≥ gene list length/2) are shown in black and red for FH and FL, respectively. Patients with positive scores in at least 50% of the gene list (corresponding to a sum of z-scores ≥ 0) are shown in grey and pink for FH and FL, respectively. **(C)** Distribution of FH and FL patients in TCGA across the Verhaak *et al* subtypes. Bar plots showing the number of FH and FL samples that fell in each Verhaak *et al* category. "Other" = patients that fell into neither category; FH&FL = patients that expressed both FH and FL genes. Mes: mesenchymal subtype; Imm: immune subtype; Pro: proliferative subtype; Diff: differentiated subtype. **(D)** ABSOLUTE (top) and ESTIMATE (bottom) algorithms applied to patients falling into the FH (black&grey), FL (red&pink) and other (blue) categories of patients using the 50% cut-off. At this less stringent cut-off the same results are obtained as shown in Figure 4C using the 75% cut-off. **(E)** Stromal content of patients falling into the FH, FL or other categories based on histopathology provided in the TCGA data set. As with the ABSOLUTE algorithm, patients falling into the "Other" category have lower stromal content than patients in the FH and FL categories. The same is true regardless of whether the 75% (top) or 50% cut-off (bottom) is applied. Related to Figure 4.

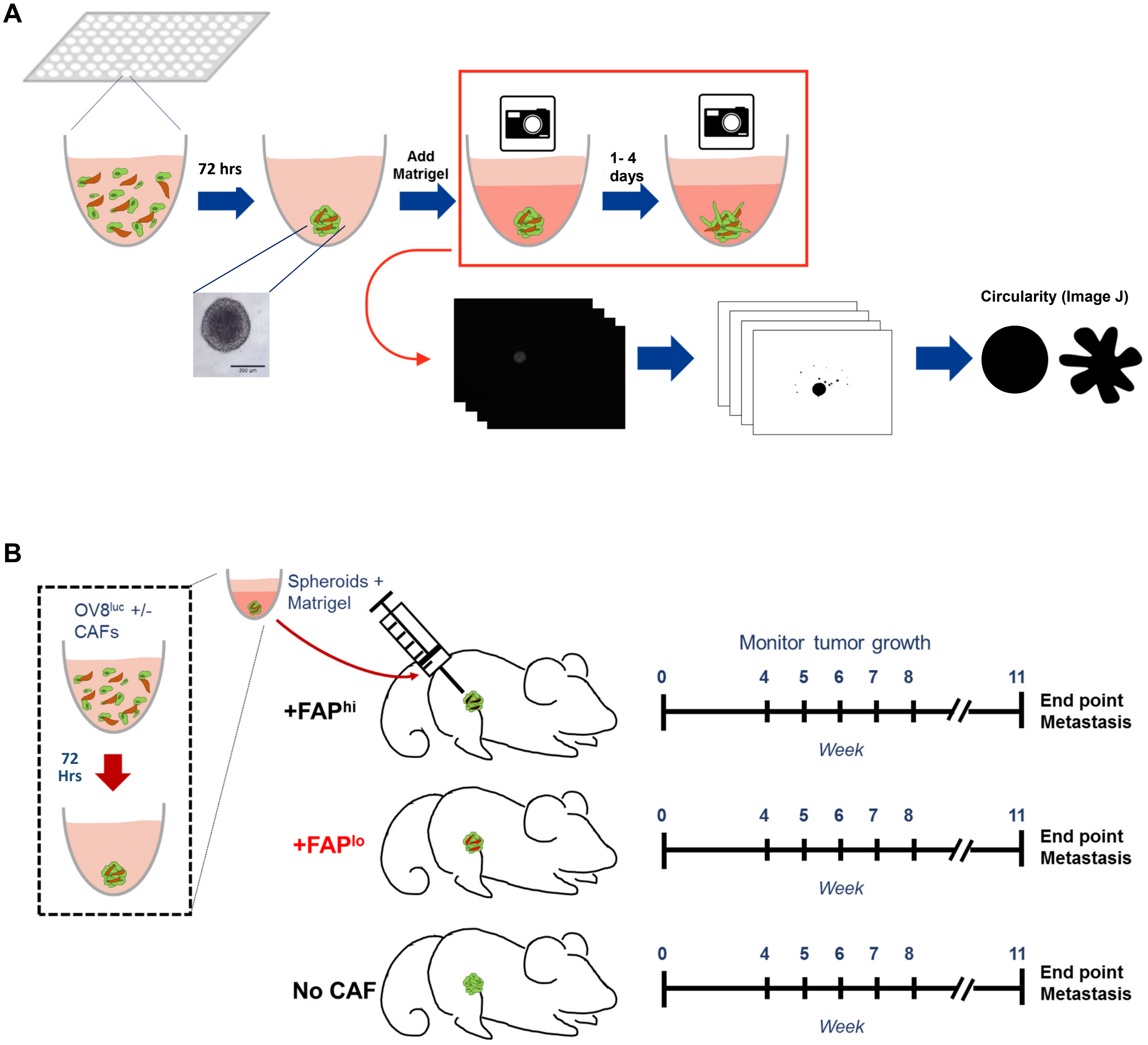


**Figure S5. Schematics for *in vitro* and *in vivo* spheroid assays. (A)** For *in vitro* invasion assays, fluorescently tagged CAFs and OVCAR8, ES2 or OV90 cells were mixed and plated into ultra-low attachment 96-well plates. Spheroids formed within 72 hours, at which time a Matrigel was added to the wells to give a final concentration of 33% v/v. Plates were gently centrifuged to center spheroids in the wells and spheroids were imaged by fluorescence microscopy to collect "time 0" images. Spheroids were then incubated for an additional 1 to 4 days, then imaged again. Image analysis was carried out using Image J software. **(B)** For *in vivo* assays spheroids were formed in the same way as in (A) using luciferase-tagged OVCAR8 cells. Once spheroids were formed, Matrigel was added to the well to give a final dilution of 50% v/v, then individual spheroids were immediately drawn up into a syringe and implanted into the mammary fat pads of NSG mice. Mice were imaged weekly using the Xenogen IVIS Imaging System 100. Related to Figures 5 and 6.

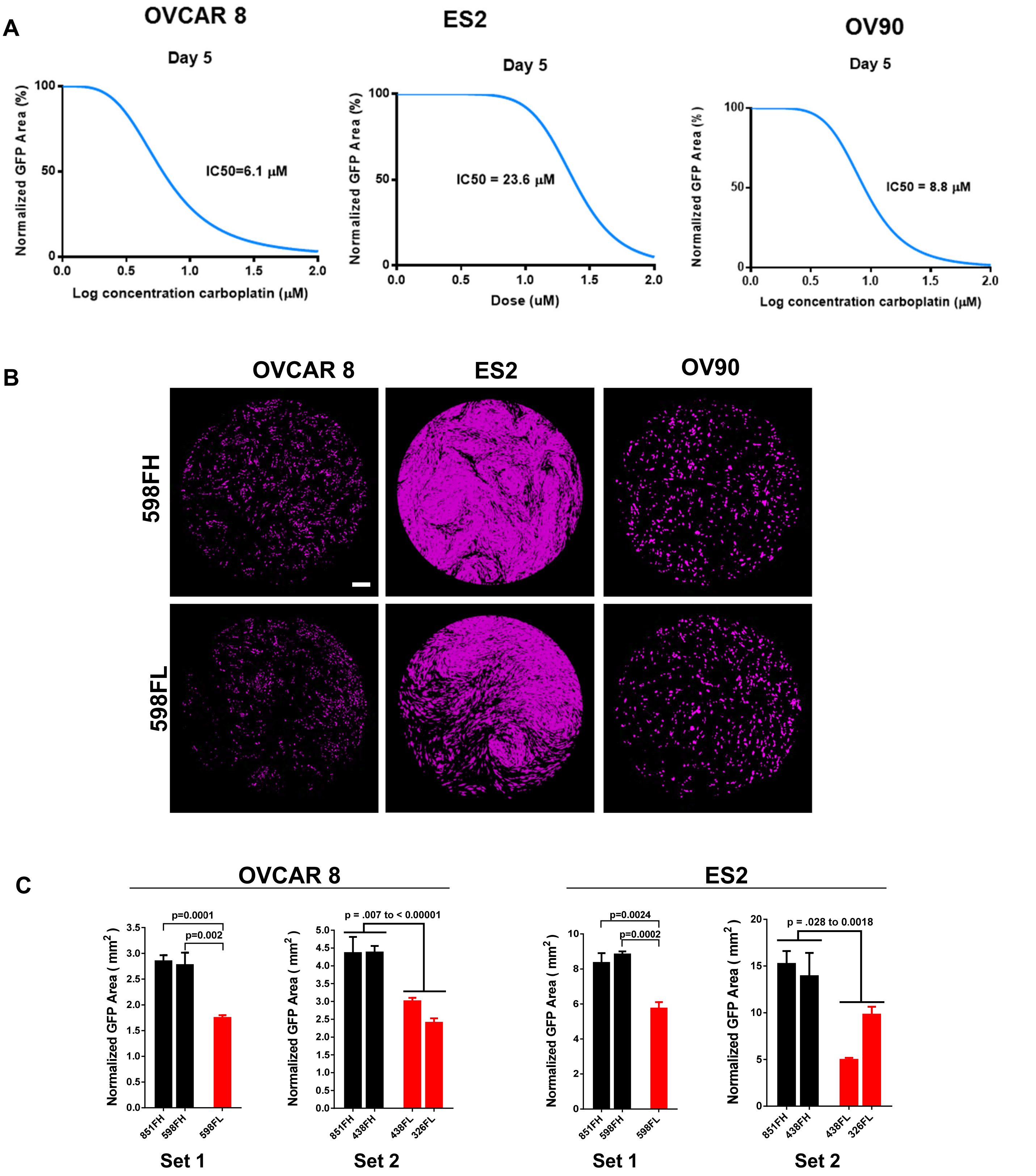


**Figure S6. Chemotherapy treatment of HGSOC cell lines. (A)** Dose response curves of OVCAR8, ES2 and OV90 cells in response to carboplatin, in adherent growth conditions. IC-50s were calculated using GraphPad software. **(B)** Representative images of OVCAR8, ES2 or OV90 cells co-cultured with 598FH (top) or 598FL (bottom) CAFs treated with 10 µM carboplatin at the 7-day time point. Pink indicates mask used to quantify GFP+ cell area. Scale bar = 100 µM. Related to Figure 5F. **(C)** Quantification of cells remaining after 7 days of treatment with 10 uM carboplatin, normalized to day 0. n=3, student's t-test.
